# The human neuronal receptor NgR1 bridges reovirus capsid proteins to initiate infection

**DOI:** 10.1101/2021.07.23.453469

**Authors:** Danica M. Sutherland, Michael Strebl, Melanie Koehler, Olivia L. Welsh, Xinzhe Yu, Liya Hu, Rita dos Santos Natividade, Jonathan J. Knowlton, Gwen M. Taylor, Rodolfo A. Moreno, Patrick Wörz, Zachary R. Lonergan, Pavithra Aravamudhan, Camila Guzman-Cardozo, David Alsteens, Zhao Wang, B. V. V. Prasad, Thilo Stehle, Terence S. Dermody

## Abstract

Human Nogo-66 receptor 1 (NgR1) is a receptor for mammalian orthoreoviruses (reoviruses), but the mechanism of virus-receptor engagement is unknown. NgR1 binds a variety of structurally dissimilar ligands in the adult central nervous system (CNS) to inhibit axon outgrowth. Disruption of ligand binding to NgR1 and subsequent signaling can improve neuron regrowth, making NgR1 an important therapeutic target for diverse conditions such as spinal crush injuries and Alzheimer disease. To elucidate how NgR1 mediates cell binding and entry of reovirus, we defined the affinity of interaction between virus and receptor, determined the structure of the virus-receptor complex, and identified residues in the receptor required for virus binding and infection. These studies revealed that NgR1 sequences in a central concave region of the molecule establish a bridge between two copies of the viral capsid protein, σ3. This unusual binding interface produces high-avidity interactions between virus and receptor and likely primes early entry steps. NgR1 sequences engaged by reovirus also are required for NgR1 binding to ligands expressed by neurons and oligodendrocytes. These studies redefine models of reovirus cell-attachment and highlight the evolution of viruses to engage multiple receptors using distinct capsid components.

## INTRODUCTION

Every virus must cross host membranes to deliver its genetic payload to the interior of the cell. To overcome this membrane barrier, viruses must successfully engage host receptors expressed on the cell surface or in endosomal compartments to mediate adhesion, internalization, and disassembly. For many viruses, accessing the cytoplasm requires more than just a “lock and key” mechanism in which a single host receptor binds a single viral capsid component. Instead, virus-receptor interactions often exist in a highly evolved landscape in which many components of a virus must engage different host molecules over time and space to orchestrate this critical step in infection. This coordinated first encounter between virus and host influences all subsequent events, including disease type and severity.

Mammalian orthoreoviruses (reoviruses) provide a highly tractable and well-established experimental system to study mechanisms of viral receptor engagement and how receptor use influences disease. Reovirus virions are nonenveloped, double-shelled particles that undergo stepwise binding to host cells and proteolytic disassembly prior to releasing a transcriptionally active viral core unit to the cell cytosol [1]. Reovirus causes age-restricted disease in many mammalian species [2–4] and readily infects humans [5–8]. In mice, reovirus disseminates from initial sites of replication in either the intestine or lung to produce strain-specific patterns of tropism in the brain and concomitant disease [9, 10]. Serotype 1 (T1) reovirus strains infect ependymal cells lining the ventricles of the brain and cause a non-lethal hydrocephalus [9], whereas serotype 3 (T3) strains infect neurons in the central nervous system (CNS) and produce a fulminant, and often lethal, encephalitis [10]. These differences in tropism and disease are mediated by the σ1 viral attachment protein and thought to be dictated by differences in receptor use [10, 11].

Most reovirus strains use a multistep binding process, engaging sequential cell-surface receptors with (*i*) low affinity, (*ii*) high affinity, and (*iii*) internalization functions to ultimately bring about uptake of reovirus virions, disassembly to form infectious subvirion particles (ISVPs) (Fig 1A), and penetration into the cytosol. Two reovirus outer-capsid proteins mediate these binding events – the σ1 viral attachment protein and the λ2 pentamer base into which σ1 embeds. Different domains of the T1 and T3 σ1 proteins bind to different sialylated carbohydrates with low affinity [12, 13], suggesting independent evolution of this binding activity for reovirus. Interactions with glycans by reovirus promote adhesion strengthening to the cell surface [14] and increase affinity for other reovirus receptors (possibly by promoting conformational changes in the viral particle) [15]. Requirements for virus binding to sialic acid differ by cell type and virus strain [12, 13, 16], and efficient glycan engagement promotes viral virulence [17–19].

**Figure 1.**
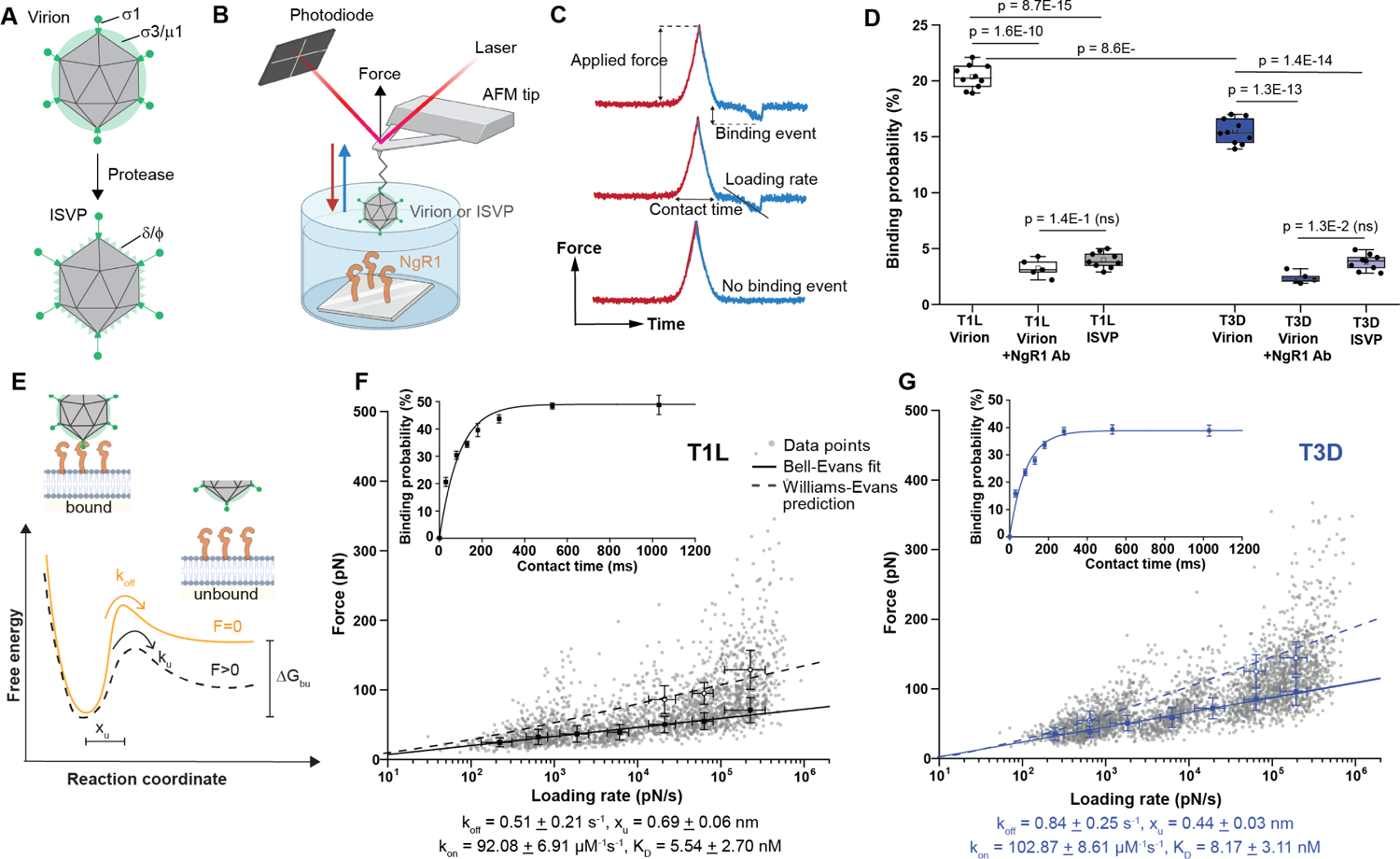
Reovirus virions bind to NgR1 on model surfaces. A Schematic of reovirus virions and infectious subvirion particles (ISVPs) showing loss of outer-capsid protein σ3, cleavage of µ1 to δ and φ fragments, and extension of the σ1 protein. B Schematic of probing reovirus particle binding to NgR1 using atomic force microscopy (AFM). Pixel-for-pixel, force-distance, curve-based AFM is used to approach and retract the sample from a tip attached to a cantilever and record interaction forces, F, over the tip-sample distance in force-distance curves. Made using BioRender. C Force-time curve from which the loading rate can be extracted from the slope of the curve immediately before bond rupture (loading rate = ΔF/Δt) (upper curve). The contact time is the interval in which the tip and surface are in constant contact (middle curve). The lower curve shows no binding events. Made using BioRender. D Box plot summarizing the binding probability from AFM studies for the conditions shown. Each data point represents the binding probability from one map acquired at a retraction speed of 1 µm/s. Mean (inset square), median (horizontal line), 25th and 75th percentiles, and highest and lowest values (whiskers) are shown. N = 10 maps examined for three 3 independent experiments. P values were determined by two-sample *t*-test. E Bell-Evans model describing a virus-receptor interaction as a two-state model (BioRender). The bound state is separated from the unbound state by a single energy barrier located at distance x_u_. k_off_ and k_on_ represent the dissociation and association rates, respectively. F, G Dynamic force spectroscopy (DFS) plot showing the distribution of average rupture forces, determined at seven distinct loading-rate ranges, quantified between NgR1 and either T1L virions (F) or T3D virions (G). Data corresponding to single interactions were fit with the Bell-Evans model (solid black or blue line, respectively), providing average k_off_ and x_u_ values. Dashed lines represent predicted binding forces for multiple simultaneous uncorrelated interactions ruptured in parallel (Williams-Evans prediction). Inserted plots: The binding probability is plotted as a function of the contact time. Least-squares fits of the data to a mono-exponential decay curve (line) provides average kinetic on-rates (k_on_) of the probed interaction. Further calculation (k_off_/k_on_) determines the K_D_. Each data point represents the binding frequency from one map acquired at a retraction speed of 1 µm/s for the different hold times. All experiments were conducted at least 3 times with independent tips and samples. Error bars indicate SD of the mean values.

The T1 and T3 σ1 proteins also bind junctional adhesion molecule-A (JAM-A), an immunoglobulin (Ig) superfamily protein, using a mechanism conserved between the serotypes [20, 21] and also conserved among many other virus-Ig superfamily receptors [22–29], in which apical viral sequences bind apical Ig receptor sequences. JAM-A expression on endothelial cells promotes reovirus viremia and hematogenous viral spread in the host [30, 31]. However, despite the high affinity of reovirus σ1 for JAM-A, this receptor does not initiate signaling required for endosomal internalization [32]. Instead, conserved sequence motifs in the viral λ2 protein promote binding to integrins, clathrin recruitment, and efficient entry [32, 33]. Binding to sialic acid and JAM-A is dispensable for reovirus replication in the intestine and brain. Thus, there are key gaps in knowledge about reovirus receptor use and function in different tissues.

Nogo-66 receptor 1 (NgR1) was identified as a reovirus receptor in an RNA interference screen using human cancer cells [34]. In contrast to JAM-A, which can bind both virions and ISVPs, NgR1 selectively binds virions [34]. ISVPs are characterized by an altered conformer of the σ1 protein and proteolytic removal of the σ3 protein, leading to the hypothesis that the virion conformer of σ1, outer-capsid protein σ3, or both engage NgR1. NgR1 is expressed in neural cells targeted by reovirus [31, 35–37] and interacts with a structurally diverse set of myelin inhibitory proteins to recruit multiple co-receptors, initiate signaling, and prevent neurite outgrowth [38–41]. There is limited structural information about how NgR1 coordinates binding to these diverse ligands and co-receptors to bring about its important functions. However, disruption of NgR1 interactions with these ligands can promote nerve growth recovery [42–45]. Thus, an enhanced understanding of the structural basis of NgR1-ligand interactions may foster development of new therapeutics.

In this study, we defined molecular interactions between reovirus and NgR1. Using a combination of biophysical, biochemical, and genetic strategies, we discovered that the reovirus outer-capsid protein σ3 serves as the viral ligand for NgR1. Indeed, two independent σ3 proteins ligate a single NgR1 molecule, a mechanism that likely promotes high-avidity interactions between virus and receptor and catalyzes early infection steps. These studies enhance an understanding of reovirus binding and entry events and also provide the first visualization of an NgR1-ligand interface.

## RESULTS

### Reovirus binds NgR1 with high avidity on receptor-coated surfaces

To determine the affinity of virus binding to NgR1 and clarify whether there are strain-specific differences in the capacity of reovirus to bind NgR1, we used force-distance-based atomic force microscopy (AFM). Virions or ISVPs of prototype reovirus strains T1L and T3D were covalently linked to an AFM probe tip, and interactions with a surface coated with purified, full-length NgR1 protein were kinetically and thermodynamically probed (Fig 1A,B). The AFM tip was cyclically approached and retracted from the NgR1-coated surface, and the force between the viral-particle-functionalized tip and the surface was monitored over time (Fig 1C; force vs. time curves). Virions of both strains displayed high binding probability to the NgR1-coated surface (Fig 1D). Importantly, pre-incubation of model surfaces with an NgR1-specific antibody abolished binding by virions, and ISVPs were incapable of NgR1 engagement (Fig 1D).

As T1L virions were ∼30% more likely to bind the NgR1-coated surface than T3D virions (Fig 1D), we speculated that sequence polymorphisms in the viral ligand might be responsible for this difference. Between these strains, the σ3 proteins share ∼97% identity, and the σ1 proteins share ∼27% identity [46]. To determine whether the T1L σ1 sequences mediate more efficient receptor engagement, we assessed the binding of reovirus strain T3SA+, which is structurally identical to T1L with the exception of a σ1 protein nearly identical to T3D. T3SA+ virions bound NgR1 comparably to T1L virions (Appendix Fig S1A), suggesting that the σ1 viral attachment protein does not mediate reovirus binding to NgR1.

We next defined the kinetic properties of the established reovirus-NgR1 bonds by probing the interaction at various loading rates (e.g., force applied over time) [47]. Fitting the data with the Bell-Evans model (Fig 1E) [48, 49] enabled us to extract the dissociation rate (k_off_) and the distance to the transition state (*x*_u_) for T1L (Fig 1F) and T3D (Fig 1G). Overall, there was a slightly higher distance to the transition state for T1L virions relative to T3D virions, indicating an increase in conformational variability following binding to NgR1, as well as a diminished dissociation rate, suggesting more stable interactions. Higher forces on dynamic force spectroscopy (DFS) plots originate from the failure of uncorrelated bonds in parallel, as predicted by the Williams-Evans model [50]. We observed good correlations between the William-Evans predictions and recorded single-molecule data. These findings suggest that reovirus virions establish multivalent interactions with NgR1.

By monitoring the influence of contact time on binding probability (Fig 1F and G; inserts), we estimated the association rate (k_on_) by assuming that the receptor-bound complex can be approximated by pseudo-fist-order kinetics [51]. Equilibrium dissociation constants (K_D_) of the studied complexes were calculated (k_off_/k_on_). All three strains tested demonstrated K_D_ values in the nM range (∼5-8 nM), indicating high-affinity interactions and further confirming the stability of the complexes established by T1L, T3D, and T3SA+ with NgR1 (insets, Fig 1F and G; Fig 3C). Similar values have been reported for other reovirus-receptor interactions [14, 33, 52]. However, the slightly lower K_D_ value for the T1L-NgR1 interaction relative to T3D-NgR1 indicates a higher affinity of reovirus T1L for NgR1. This higher affinity also is observed for the interaction of T3SA+ with NgR1 (Appendix Fig S1B, C), providing further evidence that sequence polymorphisms displayed by T1 and T3 σ1 proteins do not explain differences in biophysical interaction parameters with NgR1.

### Reovirus virions bind to NgR1 on living cells

To determine whether interactions probed on isolated NgR1 receptors are established in the context of living cells, we used Chinese hamster ovary (CHO) Lec2 cells engineered to express NgR1. CHO Lec2 cells do not express cell-surface sialic acid [53] and do not bind reovirus unless transfected with a reovirus receptor [15]. Using AFM tips functionalized with either T1L or T3D virions, we imaged confluent monolayers of co-cultured Lec2 cells (fluorescently labeled with a nuclear protein H2B-GFP and actin-mCherry) and unlabeled NgR1-expressing Lec2 cells (NgR1-Lec2) (Fig 2B and C). Fields of view were chosen in which both cell types were adjacent, serving as direct internal controls. AFM height images (Fig 2B and C; upper panels) were recorded together with corresponding adhesion images, revealing the location of specific adhesion events displayed as bright pixels (Fig 2B and C; lower panels). Lec2-NgR1 cells demonstrated a high density of adhesion events, ∼9% for T1L and ∼6% for T3D (Fig 2D), whereas Lec2 cells displayed only a sparse distribution of these events, < 2% (Fig 2D), suggesting NgR1-specific attachment of reovirus to living cells. Specificity of the probed interactions was validated by treatment with an NgR1-specific antibody prior to adsorption, leading to a significant decrease in the binding to Lec2-NgR1 cells. Similar to our findings using model surfaces, we observed a modest increase in binding to Lec2-NgR1 cells by T1L and T3SA+ relative to T3D (Appendix Fig S1D, E). Specific binding forces and corresponding loading rates were extracted from force vs. time curves recorded on Lec2-NgR1 cells and overlaid on the DFS plots obtained using NgR1-coated surfaces (Fig 2E and F; Appendix Fig S1F). Data obtained using purified receptors and living cells aligned well, confirming the relevance of results obtained using model surfaces. Collectively, these data demonstrate that reovirus binds NgR1 with high affinity.

**Figure 2.**
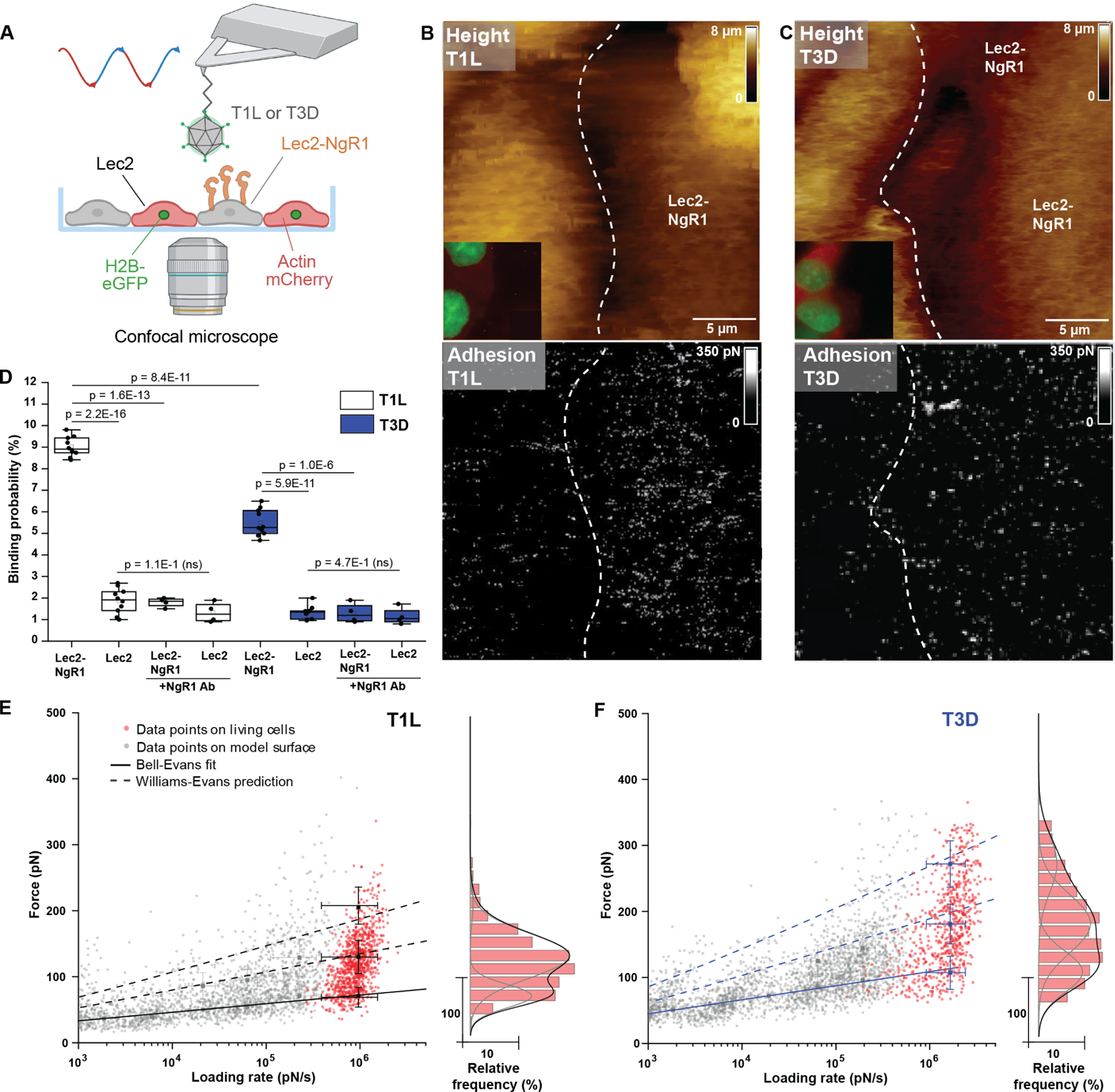
Reovirus virions bind to NgR1 expressed on living cells. A Schematic (BioRender) depicting virions probed on a confluent layer of co-cultured fluorescent Lec2 (actin-mCherry and H2B-GFP) and Lec2-NgR1 cells by combined optical microscopy and force-distance-based AFM. B, C Corresponding representative images of force-distance-based AFM topography (top panels), confocal microscopy (top panels, inset), and adhesion map (bottom panels) from probing adjacent Lec2 and Lec2-NgR1 cells with either T1L (B) or T3D (C) virions coupled to the AFM tip. Adhesion maps show interactions (white pixels) primarily with Lec2-NgR1 cells. D Box plot of the binding probability between either T1L (white) or T3D (blue) virions and Lec2 or Lec2-NgR1 cells, before and after injection of NgR1-specific antibody (Ab). Box- and-whisker plot displayed as described above. For experiments without injection of NgR1-specific antibody, the data were obtained from at least N = 10 cells from 4 independent experiments. Data for blocking experiments were obtained from at least N = 4 cells from 2 independent experiments. *P*-values were determined by two-sample *t*-test. E, F DFS plots showing the distribution of rupture forces quantified between either T1L (E) or T3D (F) virions and NgR1-overexpressing Lec2 cells (red data points). Data points in grey represent rupture forces quantified on NgR1 model surfaces (extracted from Fig 2F and G, respectively). Histograms of the force distribution observed on cells fitted with a multi-peak Gaussian distribution (N > 800 data points) are shown at the sides. Error bars indicate SD of the mean values.

### NgR1 homologs do not support reovirus infection

NgR1 displays a canonical leucine-rich-repeat (LRR) protein fold [54, 55] (Fig 3A), which is observed in many proteins, including multiple classes of pathogen sensors [56–58]. Eight tandem LRRs of NgR1 are appended on either end with non-canonical leucine-rich (LR) domains, and the C-terminus of the protein is capped by a large domain of unknown structure that anchors NgR1 to the cell membrane by a glycophosphatidylinositol (GPI) moiety (Fig 3A). NgR1 has two known homologs, NgR2 and NgR3, that have sequence similarity to NgR1 [59] and display some functional redundancy [60, 61]. To test whether NgR2 and NgR3 also serve as reovirus receptors, we transfected CHO cells with plasmids encoding NgR1, NgR2, or NgR3 and assessed expression of NgR1 and NgR2 (for which commercially available antibodies are available) (Fig 3B), reovirus binding, and infection of transfected cells. Expression of coxsackie virus and adenovirus receptor (CAR) or mock-transfection were used as negative controls. NgR1, but not NgR2 or NgR3, promoted reovirus binding (Fig 3C) and infection (Fig 3D), suggesting that NgR1 serves as a specific receptor. Since NgR3 expression could not be confirmed, these data do not exclude the possibility that NgR3 serves as a receptor for reovirus. Therefore, we assessed whether a tagged version of NgR1 retained the capacity to bind reovirus and promote infection to establish an antibody-independent method to quantify receptor expression. Myc-tagged NgR1 was detectable on the cell surface, but it promoted binding and infection much less efficiently than untagged NgR1 (Appendix Fig S2A-C), suggesting that the tag either impairs virus access to NgR1 or disrupts the structure of the receptor.

**Figure 3.**
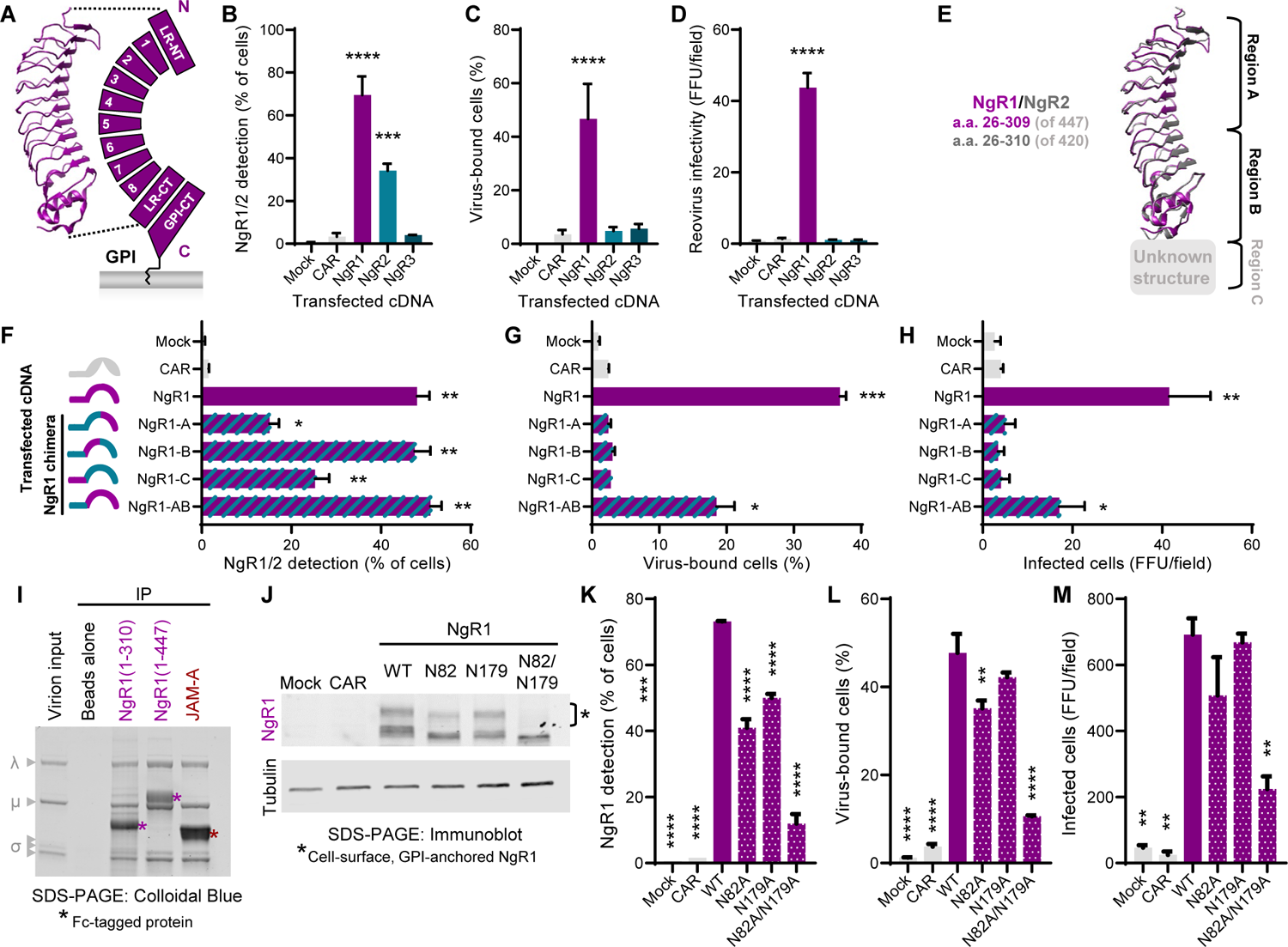
Identification of NgR1 sequences that promote reovirus binding and infectivity. A Ribbon tracing of NgR1 amino acids (a.a.) 26-310 [54] alongside a schematic of NgR1 (Leucine-rich [LR] repeat [LRR] 1-8; Glycophosphatidylinositol [GPI]). N- and C-termini are indicated. Schematic not to scale. B, C, D CHO cells were mock-transfected or transfected with the cDNAs shown and incubated for 48 h. (B) NgR1/NgR2 expression was detected on the cell surface by flow cytometry using NgR1 and NgR2 antibodies. (C) Transfected cells were incubated on ice with reovirus strain T3SA- for 1 h. Reovirus binding was detected by flow cytometry using reovirus-specific antiserum. (D) Transfected cells were incubated with T3SA-at RT for 1 h. At 24 hpi, infectivity was monitored by fluorescent focus unit (FFU) assay. Error bars indicate SD. Values that differ significantly from mock by one-way ANOVA and Dunnett’s test are indicated (***, *P* < 0.001; ****, *P* < 0.0001). E Ribbon tracings of partial NgR1 (purple) [54] and NgR2 (grey) [120] ectodomains with regions A, B, and C indicated. The structure of region C, composed primarily of the GPI-CT, has not been reported for NgR1 or NgR2. F, G, H CHO cells were mock-transfected or transfected with the cDNAs shown and incubated for 48 h. (F) NgR1 sequences were detected by flow cytometry using NgR1-specific polyclonal antiserum. (G) Reovirus binding and (H) infectivity were determined. Error bars indicate SD. Values that differ significantly from mock by student’s *t*-test are indicated (*, *P* < 0.05; **, *P* < 0.01; ***, *P* < 0.001). I Reovirus virions were incubated with protein G magnetic beads (beads alone) or beads conjugated to Fc-tagged, soluble truncated NgR1 (a.a. 26-310), full-length NgR1 (a.a. 26-447), or full-length JAM-A. Bead-bound proteins were resolved by SDS-PAGE and detected using colloidal blue staining. A representative gel from 3 independent experiments is shown. J CHO cells were mock-transfected or transfected with the cDNAs shown and incubated for 48 h. Cells were lysed, and membrane-cleared fractions were electrophoresed by SDS-PAGE and transferred to nitrocellulose. Membranes were probed for either NgR1 or tubulin and imaged. A representative gel from four independent experiments is shown. K, L, M CHO cells were mock-transfected or transfected with the cDNAs shown and incubated for 48 h. (J) NgR1 expression, (K) reovirus binding, or (L) reovirus infectivity were determined. Error bars indicate SD. Values that differ significantly from mock by one-way ANOVA and Dunnett’s test are indicated (*, *P* < 0.05; **, *P* < 0.01; ***, *P* < 0.001).

### NgR1 amino acids 26-310 are necessary and sufficient to bind reovirus

The repetitive solenoid structure of NgR1 makes it particularly amenable to deletion, replacement, or addition of LRR domains [55, 60–62]. As a first step to define domains of NgR1 required for reovirus binding, we engineered a cDNA panel encoding sequential deletions of NgR1 domains (ΔLR-NT, ΔLRR1-2, ΔLRR3-4, ΔLRR5-6, ΔLRR7-8, ΔLR-CT, and ΔGPI-CT) and tested the capacity of these constructs to bind virus and allow infection. All constructs were detected on the cell surface, but none promoted reovirus binding or infectivity (data not shown), indicating that the reovirus binding epitope was disrupted in each case.

As a complementary gain-of-function approach, we engineered chimeric constructs in which NgR2 sequences were exchanged with NgR1 sequences to identify NgR1 sequences required to confer receptor activity. Although all of the chimeric proteins were detected on the cell surface (Fig 3F), only the construct expressing the full complement of NgR1 LR sequences appended to the anchor domain of NgR2 (NgR1-AB) promoted reovirus binding (Fig 3G) and, to a modest extent, infection (Fig 3H). These data suggest that LR domains of NgR1 are required for binding and, furthermore, suggest that the membrane-proximal GPI-CT anchor domain is dispensable for interactions with reovirus. To determine whether the NgR1 GPI-CT domain contributes to virus binding, we tested either full-length NgR1 or truncated NgR1 expressing only LR sequences (NgR1 a.a. 1-310) for the capacity to bind reovirus virions. Corroborating our experiments with NgR1/NgR2 chimeras, truncated NgR1 protein efficiently precipitated reovirus virions (Fig 3I).

### N-glycosylation of NgR1 LRR domains is dispensable for reovirus binding

As many reovirus strains interact with sialic acid, we tested whether NgR1 glycosylation contributes to reovirus binding. NgR1 is glycosylated at asparagine residues 82 and 179, which are located in the central LRR region, as well as several N- and O-linked sites in the GPI-CT domain [54, 63]. To determine whether NgR1 glycosylation is required for reovirus receptor activity, we exchanged N82 and N179 with alanine either singly or in combination and assessed reovirus binding and infectivity using receptor-transfected CHO cells. Immunoblotting of transfected cell lysates revealed predicted shifts in the electrophoretic mobility of the mutant NgR1 constructs (approximately 1-2 kDa per glycosylation site) (Fig 3J), suggesting that the mutants lacked the targeted glycosylation sites. Protein bands corresponding to the mature form of NgR1 (Appendix Fig S3) were relatively absent for the N82/N179 double mutant (Fig 3J and S4; asterisk), indicating that this mutant has impaired conformation or stability. While cell-surface levels of NgR1 single-glycosylation mutants N82A and N179A were less than those of wild-type NgR1 (Fig 3J and K), both promote infection comparable to wild-type NgR1 (Fig 3M). Interestingly, the small amount of NgR1-N82A/N179A detectable on the cell surface (Fig 3J and K) appears to contribute to modest virus binding (Fig 3L) and infection (Fig 3M). Thus, reovirus can use NgR1 as a receptor independent of glycosylation sites in the central LR protein region.

### NgR1 engages the σ3 outer-capsid protein

NgR1 promotes binding and infection of reovirus virions but not ISVPs [34]. This particle-specific binding capacity suggests that either the more compact virion conformer of σ1, σ3, or some combination of these capsid components serves as the viral ligand for NgR1. To identify the viral NgR1 ligand, we determined whether soluble, purified NgR1 can interact with isolated σ1 protein or σ3 protein. The virion particle contains 12 σ1 trimers at the virion icosahedral fivefold axes and 200 σ3-μ1 heterohexamers, with each composed of three copies of σ3 resting on a pedestal of three copies of μ1 protein (σ3_3_μ1_3_) (Fig 4A). Co-expression and purification of σ3 and μ1 produces heterohexameric sub-assemblies, in which σ3 is displayed in its native virion state bound to μ1 (Fig 4A and B) [64]. Using NgR1-Fc as an affinity ligand in a protein G bead-based binding assay, we observed NgR1-Fc binding to σ3_3_μ1_3_ complexes (Fig 4C, D) but not to σ1 (data not shown). Importantly, this binding is highly specific, as the unrelated CAR protein was unable to precipitate σ3_3_μ1_3_.

**Figure 4.**
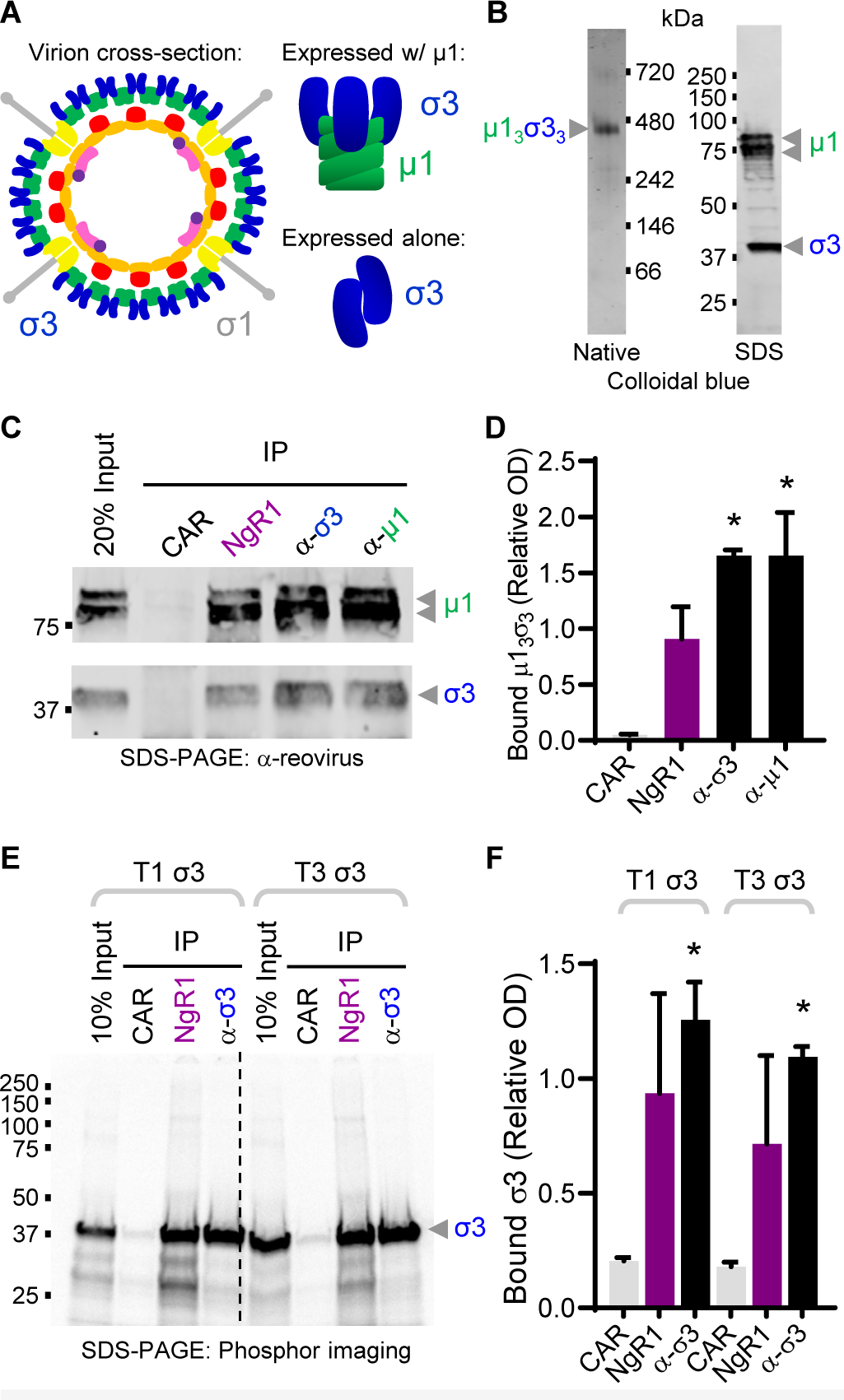
Reovirus outer-capsid protein σ3 is the viral ligand for NgR1. A Schematic of reovirus capsid components. B Purified σ3_3_μ1_3_ electrophoresed by native page (left) or SDS-PAGE (right) and stained by colloidal blue. C, D Soluble Fc-tagged CAR or NgR1, or monoclonal antibodies 10C1 (σ3-specific) or 8H6 (μ1-specific) were immobilized onto protein G beads. Protein-conjugated beads were incubated with purified σ3_3_μ1_3_ heterohexamer. Beads were washed, boiled, and released proteins were electrophoresed by SDS-PAGE and visualized using reovirus-specific antiserum. A representative gel (C) and quantification of two independent experiments (D) are shown. SEM are shown. Values that differ significantly from CAR by *t*-test are indicated (*, *P* < 0.05). E, F Soluble Fc-tagged CAR or NgR1 or 10C1 (σ3-specific) monoclonal antibody were incubated with rabbit reticulocyte lysates expressing radiolabeled T1L or T3D σ3. Beads were washed, boiled, and released proteins electrophoresed by SDS-PAGE. Immunoprecipitated protein was visualized by phosphor imaging. A representative gel (E) and quantification of 3 independent experiments (F) are shown. Values that differ significantly from CAR by one-way ANOVA and Dunnett’s test are indicated (*, *P* < 0.05).

When expressed without μ1, σ3 forms dimers (Fig 4A) [65], shielding some surfaces that would otherwise be exposed in virion-associated conformers of σ3. To determine whether σ3 alone is capable of binding NgR1, we translated σ3 in mRNA-depleted rabbit reticulocyte lysates in the presence of ^35^S methionine. Radiolabeled σ3 was then incubated with CAR-Fc, NgR1-Fc, or a σ3-specific antibody, and Fc-expressing proteins were precipitated using protein G beads. Remarkably, isolated σ3 was capable of specifically binding NgR1 (Fig 4E and F). Moreover, we did not observe major differences in NgR1 binding by T1L and T3D σ3, consistent with previous findings (Fig 1) [34]. Collectively, these data reveal that σ3 is the viral ligand for NgR1.

### Cryo-EM reconstructions reveal an unusual binding mode of NgR1 to bridge two σ3 capsid proteins

To elucidate the structure of NgR1 bound to σ3, we conducted cryo-electron microscopy (cryo-EM) analyses of NgR1 bound to whole virions. Purified virions were incubated alone or with soluble NgR1 protein, and samples were imaged by cryo-EM. Virus particles incubated with buffer alone displayed high-contrast, regular margins (Fig 5A), whereas those incubated with NgR1 displayed extended, hazy margins (Fig 5B), suggestive of NgR1 binding. We averaged several hundred particles of virions alone or virions in complex with NgR1 to build 3D reconstructions at final resolutions of 7.2 Å and 8.9 Å, respectively. Icosahedral symmetry of the virion, which was applied during reconstructions and anticipated based on available structures of reovirus virions [66, 67], is clearly evident in both reconstructions by the formation of 2-fold, 3-fold, and 5-fold axes of symmetry (Fig 5C to 5F). Importantly, additional density was detected at the surface of virions complexed with NgR1, which can be observed in cross-sections of the reconstructions (Fig 5D, arrowheads) and the three-dimensional overview in which additional density appears in a star-like pattern overlying σ3 protein (Fig 5F). Consistent with co-precipitation assays (Fig 4E and F), these data demonstrate that NgR1 binds to σ3.

**Figure 5.**
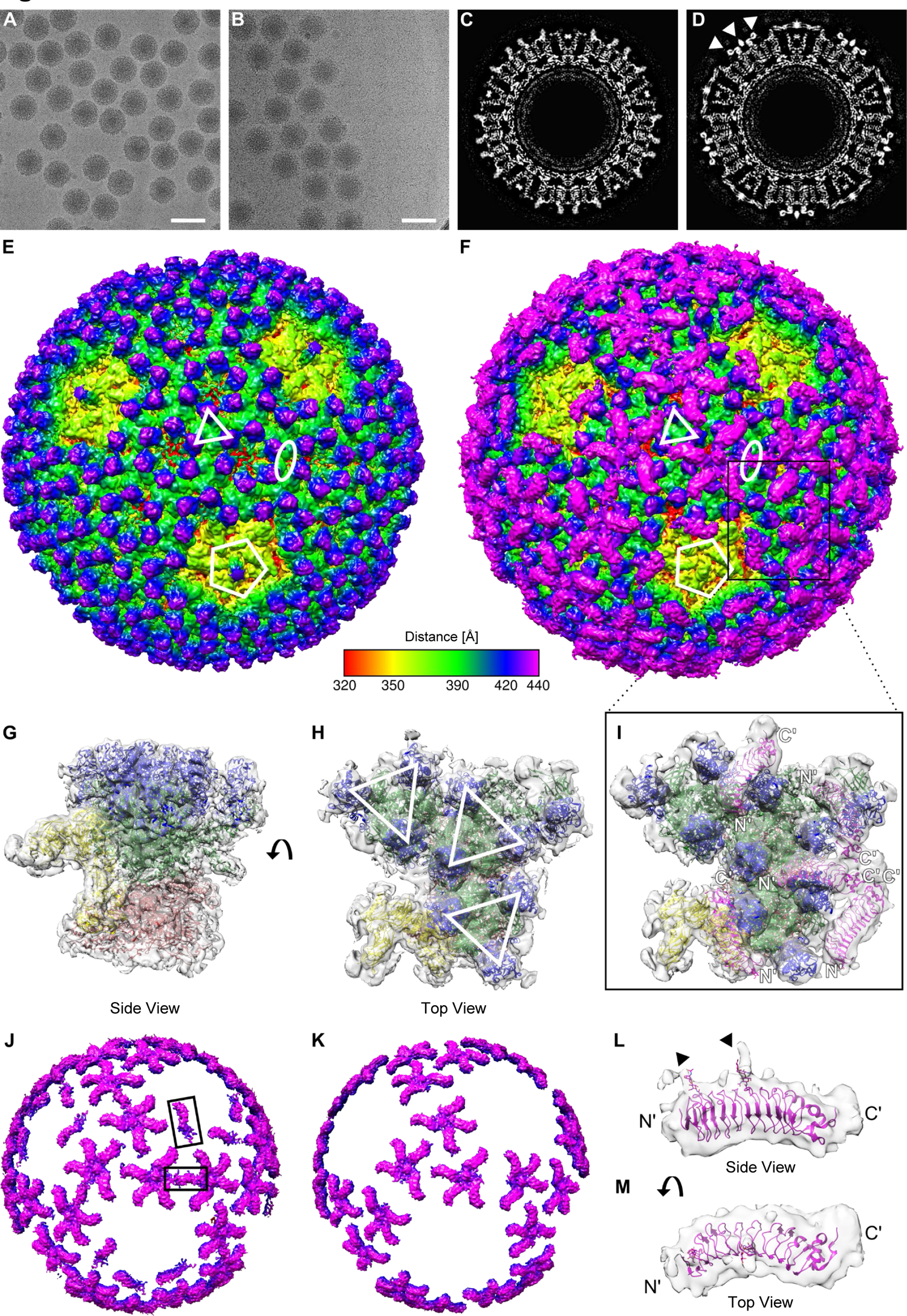
Cryo-EM reconstructions of NgR1 bound to reovirus virions reveals the interaction arrangement. A, B Representative cryo-electron micrographs of (A) reovirus virions or (B) reovirus virions complexed with NgR1. Scale bar, 100 nm. C, D Central slices of the 3D cryo-electron reconstruction of (C) reovirus or (D) reovirus complexed with NgR1. White arrowheads indicate NgR1 density. E, F Overview of 3D reconstructions for (E) reovirus or (F) reovirus complexed with NgR1 at resolutions of 7.2 Å and 8.9 Å, respectively. Maps are colored by distance from the core (320 Å −440 Å). Representative 2-fold, 3-fold, and 5-fold symmetry axes of the icosahedral capsid are indicated by white symbols. G, H, I Asymmetric units for reovirus (G, side view; H, top view) or reovirus-NgR1 (I, top view). Ribbon tracings of reovirus proteins (σ3-blue, μ1-green, λ2-yellow, λ3-red, and σ2-red) and NgR1 (magenta) were placed into the 3D reconstructions (grey translucent surface representation) of virions (G,H) or the virion-NgR1 complex (I). (H) Individual σ3_3_μ1_3_ heterohexamers are indicated with white triangles. (I) N- and C-termini for docked NgR1 molecules are labeled in white. J, K Difference density maps of the reovirus virion reconstruction subtracted from the reovirus-NgR1 reconstruction at sigma level 1.5 (J) or 2.0 (K). Major additional NgR1 features are indicated by black rectangles. L, M Side and top views of NgR1 ribbon tracing placed into the difference density map (sigma level = 1.15). Glycosylations are shown in stick format, and densities attributable to glycosylation are indicated by black arrows.

To obtain molecular detail about the σ3-NgR1 binding surface, we docked available protein structures into the cryo-EM density maps (Fig 5G to I). The asymmetric reovirus unit, which can be tiled 60 times to build a particle *in silica*, consists of one copy each of λ2, σ2, and λ1, and ten copies each of μ1 and σ3. These ten molecules of μ1 and σ3 compose three full heterohexamers (Fig 5H, indicated by white triangles) and one third of a fourth heterohexamer, which is located at a three-fold symmetry axis. The flexibility, trimeric nature, and incomplete occupancy of σ1 prevent resolution of this protein in most particle-averaged reconstructions, but remnants of the fiber base embedded into the λ2 pentamer are occasionally detected (Fig 5E, purple nub above yellow) [67]. Within the asymmetric unit of the reovirus-NgR1 reconstructions, five NgR1 molecules fit well within the remaining cryo-EM features (Fig 5I) and display binding modes comparable to each other (Appendix Fig S5). Four of these NgR1 molecules are embedded between two σ3 monomers of neighboring heterohexamers, while the fifth is located adjacent to the five-fold vertex and thus is only flanked by one σ3 monomer. This structure reveals that two binding faces can be used by both NgR1 and σ3. Moreover, NgR1 density is distant from μ1, confirming a specific interaction of the receptor with σ3.

To define additional features of the reovirus-NgR1 interaction, we subtracted the density of the reovirus virion reconstruction from that of the reovirus-NgR1 reconstruction and evaluated the resulting difference density map at different threshold levels (Fig 5J and K). NgR1 can be unambiguously placed in all five of the binding sites in the asymmetric unit (Appendix Fig S5). We also detected another feature in the difference density map located at the two-fold symmetry axes of the virion surface (Fig 5F and J). This density is not shaped like a single NgR1 molecule but rather like an average of two NgR1 molecules, each bound in opposing directions. Therefore, accurate placement of the NgR1 coordinates into this density was not possible. However, the location of the density between σ3 monomers of adjacent hexamers is consistent with the NgR1 binding mode detected nearby on the particle. Increasing the threshold level of the difference density map from 1.5 to 2.0 removes this feature, along with the NgR1 moieties observed at the five-fold vertex (Fig 5J and K). This signal loss suggests weaker NgR1 density at these positions, which might be caused by decreased binding affinity to these sites.

The correct orientation of NgR1 is confirmed by the glycosylation signatures at N82 and N179, which are observed at a slightly lower threshold level (1.15) (Fig 5L and M). These glycan chains face away from the virion surface, which likely explains why they do not contribute to reovirus binding and infection (Fig 3L and M). This orientation allows the His tag appended to NgR1, which replaces the GPI-CT of full-length NgR1, to accumulate at the center of four-to-five bound NgR1 molecules (Fig 5I and Fig 6A), suggesting a role for NgR1 oligomerization in binding to reovirus.

**Figure 6.**
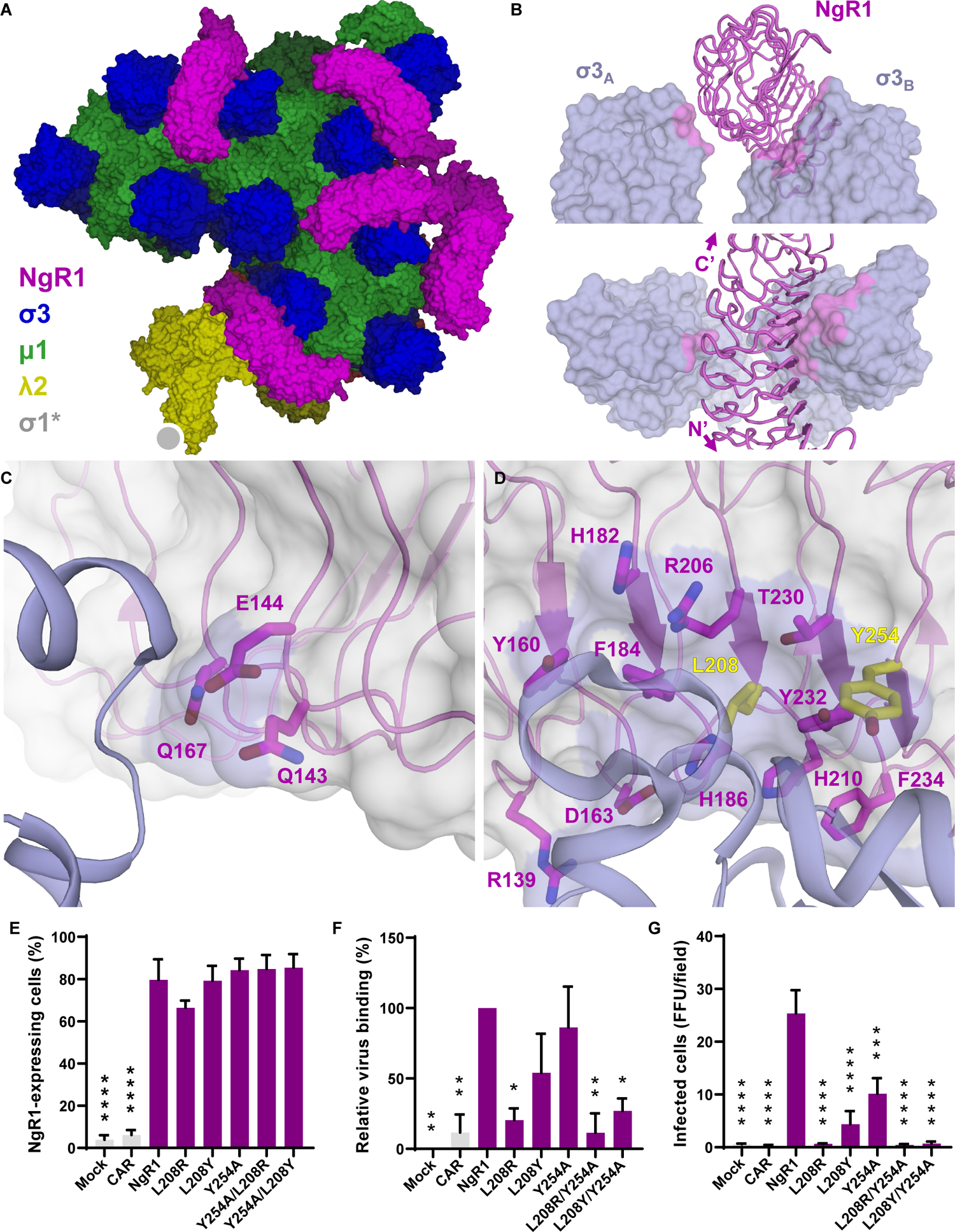
NgR1 residues required for interaction with reovirus identified by structure-guided mutagenesis. A Surface representation of the modelled asymmetric unit of a reovirus virion in complex with NgR1 (extracted from the Figure 5 molecular docking). NgR1 is colored in magenta, σ3 in blue, μ1 in green, λ1 in yellow, and σ2 as well as λ1 in red. The approximate location at which σ1 imbeds into λ2 is depicted by a grey circle. B Interactions of NgR1 with two σ3 monomers (labelled σ3_A_ and σ3_B_) from different heterohexamers are depicted in a side view (top panel) and top view (bottom panel). NgR1 is shown as a ribbon tracing (magenta) and σ3 as a surface representation (blue). The σ3 surface < 5 Å from the NgR1 protein model is colored in magenta. C, D Magnified views of the σ3_A_ (C) and σ3_B_ (D) interaction sites. NgR1 amino acids in proximity to σ3 are shown in stick representation. Residues with confirmed critical functions in NgR1 binding, L208 and Y254, are colored in yellow. E, F, G CHO cells were mock-transfected or transfected with the cDNAs shown and incubated for 48 h. (E) NgR1 expression, (F) reovirus binding, or (G) reovirus infectivity were determined. Error bars indicate SD. Values that differ significantly from mock by one-way ANOVA and Dunnett’s test are indicated (*, *P* < 0.05; **, *P* < 0.01; ***, *P* < 0.001; ***, *P* < 0.001).

By identifying NgR1 residues within 5 Å of σ3 (Fig 6B), we predict two important interaction interfaces, a smaller surface on the convex surface of NgR1 and a significantly larger one on the more-conserved concave surface of NgR1 (Fig 6B). The smaller surface consists mostly of glutamines and glutamates that could potentially form electrostatic or polar interactions (Fig 6C). The larger surface is defined by a broad patch of aromatic residues with a central cavity filled by L208 and two prominent residues, R139 and Y254, adjacent to the interface (Fig 6D). This binding surface provides possibilities for charged, polar, or hydrophobic interactions.

### NgR1 residues required for σ3 binding reside in a pocket

To identify NgR1 residues required for binding to σ3, we used the reovirus-NgR1 structure to rationalize mutations at key sites in the interaction interface. We exchanged L208 with arginine or tyrosine and Y254 to alanine, either alone or in combination with the L208 mutants. The mutant NgR1 constructs were detectable on the CHO cell surface (Fig 6E), but all displayed impaired capacity to bind reovirus (Fig 6F) and failed to efficiently promote infection (Fig 6G). Combination of mutations at L208 and Y254 had synergistic effects in diminishing reovirus binding and infectivity (Fig 6F and G). These data demonstrate that NgR1 residues L208 and Y254, which reside at the NgR1-σ3 interface, are required for reovirus binding and infection and provide clues about the binding mechanism.

## DISCUSSION

All viruses must specifically engage host cell-surface molecules to achieve reliable membrane apposition and cell entry. In this study, we evaluated biophysical parameters of reovirus binding to NgR1 and functional outcomes of this interaction. Collectively, our results reveal high-affinity interactions between reovirus virions and NgR1 as well as an unusual binding mode for virus-receptor interactions in which NgR1 rests between two adjacent capsid proteins. These studies expand our understanding of the reovirus capsid components used to bind cells and highlight the complex and multimodal strategy in which reovirus engages several receptors using different capsid components to promote efficient infection.

We chose cryo-EM to resolve NgR1 bound to whole reovirus particles. Similar approaches have resolved structures of other virus-receptor interactions [25, 26, 68, 69]. Due to the multiple possibilities of binding (two faces of NgR1 and two faces of σ3), we thought it unlikely that the purified interactants would form stable, regular complexes required for a crystallographic determination of the structure. Ongoing advances in cryo-EM methods and analyses [70] now allow structural determination at near atomic-level resolution. While the structures we report here do not achieve atomic-level resolution, combining the 3D reconstructions we obtained with molecular docking and functional analyses provides confidence in how NgR1 binds the reovirus capsid.

In addition to the newly discovered function of reovirus σ3 in NgR1 binding, σ3 has several other important activities. During replication, σ3 binds dsRNA [71], prevents activation of protein kinase R (PKR) [72, 73], and suppresses stress granule formation [74], thereby promoting efficient viral replication [74, 75]. As a component of incoming virion particles, σ3 is thought to serve as a protector protein for the μ1 protein, which functions in membrane penetration. The σ3 protein must be cleaved and removed by cathepsin or intestinal proteases to expose and prime μ1 to penetrate membranes [1]. How reovirus interactions with NgR1 alter particle stability, σ3 cleavage or uncoating, and early entry steps remains to be determined. As NgR1 is GPI-anchored and has no cytoplasmic domain, intracellular signaling following NgR1 ligand binding is mediated by NgR1 co-receptors on neurons. Several co-receptors have been described [76–79], but it is not known whether these function in reovirus binding or infection.

In the CNS, NgR1 binds a structurally diverse set of myelin-associated inhibitory ligands, e.g., Nogo-A [38], oligodendrocyte myelin-associated glycoprotein (OMgp) [41], and myelin-associated glycoprotein (MAG) [39]. While structural data are available for individual components [80–82], it is not known how these myelin-associated inhibitory molecules interact with NgR1 in complex. To our knowledge, we present the first structure of NgR1 bound to a non-antibody ligand. Reovirus engages N-terminal, LR sequences of NgR1 at two distinct sites (Fig 6B). A smaller patch on the back and side of NgR1 (Fig 6B and C) coordinates with a broad patch of residues on the concave surface of NgR1 (Fig 6B and D) to imbed NgR1 between two σ3 proteins (Fig 5F and 6A and B). Mutagenesis studies confirm the requirement of two key residues at the center and periphery of the larger NgR1-σ3_B_ surface in reovirus binding and infectivity (Fig 6E-F). Interestingly, many native binding partners of NgR1 similarly require N-terminal LR sequences (a.a. 26-310) for binding [61, 62], suggesting that the reovirus binding sites on NgR1 overlap with those of the myelin-associated inhibitory ligands engaged in the CNS.

A key remaining question is why do reovirus, myelin-associated inhibitory ligands, and even some NgR1 co-receptors [62] target and even compete [39] for these central LR sequences? One possible explanation is the relatively high conservation of sequences on the concave NgR1 LR surface [54, 55]. As NgR1 serves important functions in both the developing and mature CNS, maintenance of ligand interactions is likely to foster sequence conservation at these interfaces. In line with this idea, single nucleotide polymorphisms of NgR1 in human populations have been associated with schizophrenia [83–85] and other neurologic disorders. Indeed, many viruses, including reovirus [21], benefit from engaging protein surfaces that are required for critical interactions with other host proteins, and thus are strictly conserved.

An alternative possibility is that the residues engaged by reovirus and other NgR1 ligands are those most readily displayed by the native structure of NgR1 on cells. While many viruses bind the most apical surface of an extended receptor [22–29], there are few examples of virus binding to lateral receptor surfaces [86, 87]. The precise orientation of NgR1 on the cell surface is not known, however, some hypothesize that the concave NgR1 surface faces toward ligands on neighboring cells. This hypothesis is consistent with the identification of a hydrophobic pocket at the base of the convex surface of NgR1 [54], which may interact with the unresolved GPI-CT structure to allow display of the expansive concave surface for efficient ligand interactions. This NgR1 arrangement could explain the unusual *en face* arrangement of NgR1 that we observe on virions (Fig 5F and 6A). Additionally, clustering of NgR1 C-termini at the center of σ3_3_μ1_3_ heterohexamers (Fig 5F) suggests that NgR1 interacts with itself in a complex to bind reovirus, which could promote higher avidity binding. There is evidence that NgR1 forms homophilic interactions [40, 88] and that NgR1 is not broadly distributed in the membrane but instead clusters in lipid-rich microdomains [89]. However, the nature of these interactions and their effect on reovirus binding is not clear. Additional studies of the GPI-CT structure and its involvement in NgR1 conformation and function will be required to answer these questions.

Reovirus interacts with several cellular receptors using distinct capsid components. The fiber-like σ1 viral attachment protein engages sialylated glycans and JAM-A [12, 13, 20, 21], while the λ2 pentameric protein engages β1 integrin [33]. We discovered that NgR1 serves as a receptor of reovirus σ3, which expands our understanding of reovirus interactions with host cells. Interestingly, our studies also reveal yet another parallel of reovirus structure and function to adenoviruses [90, 91], which engage CAR/sialylated glycans, integrins, and CD46 receptors using fiber [25], penton [92], and hexon [87] structural proteins, respectively. These shared similarities demonstrate how some viruses have evolved the use of all surface-exposed capsid components as receptor-binding proteins and highlight the critical importance of receptor engagement.

Reovirus tropism for neurons and its capacity for neurovirulence are exquisitely dictated by sequences in the T3 σ1 head domain [11]. Consistent with previous findings [34], we observe no overt preference for NgR1 to engage T3 sequences (Fig 4E and F). In fact, NgR1 may bind T1 σ3 with slightly higher affinity relative to T3 σ3 (Fig 1F and G; Appendix Fig S1C). These data suggest that other receptors are engaged by reovirus *in vivo* to cause encephalitis. However, not all neurotropic orthoreoviruses target neurons in this σ1-dependent fashion. Baboon reovirus (BRV), which causes encephalitis in baboons [93, 94], lacks a σ1-like attachment protein but otherwise resembles reovirus structurally [95]. BRV and other relatives of the strains tested here should be evaluated to understand the viral and host species determinants that allow reovirus to bind NgR1 and how these interactions influence target cell selection and disease. As NgR1 was first identified as a receptor for reovirus from a screen using human cancer cells [34] and reovirus is being evaluated as an oncolytic in humans [96, 97], these studies also may improve the targeting of oncolytic therapeutics. Moreover, findings presented here may inform an understanding of how NgR1 engages structurally diverse ligands, and furthermore, how these interactions influence functions of NgR1 in shaping the CNS.

## MATERIALS AND METHODS

### Cells

Spinner-adapted L929 fibroblasts were maintained either in suspension culture or adherent in Joklik’s minimal essential medium (JMEM) supplemented to contain 5% fetal bovine serum (FBS), 2 mM L-glutamine, 100 U/mL penicillin, 100 µg/mL streptomycin, and 0.25 µg/mL amphotericin B. CHO Lec2 cells (ATCC; CRL-1736) were transduced with lentiviruses expressing actin-mCherry and H2B-GFP. Cells expressing both GFP and mCherry were sorted by flow cytometry and further propagated (Lec2) [15]. Independent CHO Lec2 cells were transduced with lentivirus encoding human NgR1 (NM_023004), and NgR1-expressing cells were sorted by flow cytometry and further propagated (Lec2-NgR1). Lec2 and Lec2-NgR1 cells were maintained in MEM α nucleosides medium (Gibco) supplemented to contain 10% FBS, 100 units/mL penicillin, 100 µg/mL streptomycin, and 2 mM L-glutamine. CHO cells (ATCC, CCL-61) were maintained in Ham’s F12 medium supplemented to contain 10% FBS, 100 U/ml penicillin, 100 µg/ml streptomycin, and 0.25 mg/ml amphotericin B. Freestyle™ 293-F cells (Thermo Fisher Scientific) were maintained in FreeStyle™ 293 Expression Medium as recommended (Thermo). All mammalian cells were cultivated at 37°C in a humidified atmosphere with 5% CO2. High Five™ insect cells (Thermo Fisher Scientific) were maintained in rotating suspension cultures as recommended (Thermo).

### Antibodies

Reovirus polyclonal antiserum was collected from rabbits immunized and boosted with reovirus strain T1L or T3D. Sera from T1L- and T3D-inoculated rabbits were mixed 1:1 (vol:vol) and adsorbed on L929 cells to deplete non-specific antibodies. The following antibodies were used in specified assays at the indicated dilutions: anti-reovirus rabbit serum (FFU assay – 1:1000; flow cytometry – 1:20,000; immunoblot – 1:1000); anti-NgR1 goat polyclonal IgG (R&D Systems; AF1208) (AFM blockade – 100 μg/mL; flow cytometry – 0.2 μg/mL; immunoblot – 0.2 μg/mL); anti-NgR2 goat polyclonal antibody (R&D Systems; AF2776) (flow cytometry – 0.1 μg/mL); anti-α-tubulin mouse monoclonal antibody (Cell Signaling Technology; clone DM1A) (immunoblot – 1:2000); anti-myc mouse monoclonal antibody (Cell Signaling Technology; clone 9B11) (flow cytometry – 1:1000. Anti-σ3 (10C1) or anti-μ1 (8H6) mouse monoclonal antibodies [98] were used for immunoprecipitations as described below.

### Viruses

Reovirus strains T1L and T3D are prototype serotype 1 and serotype 3 viruses that were originally isolated from infected children [99]. Strains T3SA+ and T3SA-are engineered recombinant strains [11] that express nine genes of T1L (including the S4 gene that encodes σ3) and the S1 gene of either T3-clone 44-MA or T3-clone-44 [100], respectively. T3SA+ and T3SA-virions differ by a single amino acid polymorphism (σ1-P204L) that abrogates sialic acid binding capacity of the T3SA-σ1 protein [100]. All viruses were recovered using plasmid-based reverse genetics [101, 102]. Virus was purified from infected L929 cells by cesium chloride gradient centrifugation [103]. Viral titers were determined by particle number (estimated by spectral absorbance at 260 nm [1 OD^260^ = 2.1 ˣ 10^12^ particles/mL]) or plaque assay in the absence of exogenous proteases and presence of FBS [103].

### Expression plasmids

All studies here used human Nogo receptor sequences, following a HeLa cell screen and validation of hNgR1 as a reovirus receptor [34]. Constructs used in this study are indicated in the table below, with amino acids (a.a.) for NgR1, NgR2, and NgR3 annotated. Nogo receptor sequences at the N- and C-termini of the protein are removed by signal peptide cleavage and addition of a GPI anchor, respectively. Therefore, annotated sequences reflect what is encoded by the cDNA, and not the mature protein. Human cDNA of CAR and JAM-A in pcDNA3.1 have been described [52]. Human cDNA (Origene) of NgR1 (NM_023004), and NgR2 (NM_178570), and NgR3 (NM_178568) were sub-cloned into the pcDNA3.1+ expression plasmid using sticky-end mutagenesis and custom primers. Myc-hNgR1 in the pSecTag2 expression vector was provided by Dr. Stephen M. Strittmatter (Yale University). The S4 gene ORF (σ3-encoding sequence) of reovirus strain T1L (GenBank Accession: M13139.1) or T3D (GenBank Accession: HM159622.1) were sub-cloned from pT7 [101] into the pcDNA3.1+ expression plasmid using Gibson assembly and custom primers.

| Amino Acids of Human Nogo Receptors |  |  |  |  |  |
| --- | --- | --- | --- | --- | --- |
|  | NgR1 | NgR2 | NgR3 | Plasmid | Notes |
| *Wild-type* |  |  |  |  |  |
| NgR1 | 1-473* |  |  | pcDNA3.1+ |  |
| NgR2 |  | 1-420** |  | pcDNA3.1+ |  |
| NgR3 |  |  | 1-441*** | pcDNA3.1+ |  |
| NgR1-myc | 1-473 |  |  | pSecTag2 | N-term Myc |
| NgR1-His | 1-310 |  |  | pcDNA3.1+ | C-term 7X His |
| NgR1(1-447)-Fc | 1-447 |  |  | pcDNA3.1+ | C-term Fc |
| NgR1(1-310)-Fc | 1-310 |  |  | pcDNA3.1+ | C-term Fc |
| *Glycosylation mutant* |  |  |  |  |  |
| NgR1 N82A | 1-473 |  |  | pcDNA3.1+ | N82A |
| NgR1 N179A | 1-473 |  |  | pcDNA3.1+ | N179A |
| NgR1 N82A/N179A | 1-473 |  |  | pcDNA3.1+ | N82A/N179A |
| *Chimeric NgR1/NgR2* |  |  |  |  |  |
| NgR1-A | 1-154 | 156-420 |  | pcDNA3.1+ |  |
| NgR1-B | 155-310 | 1-155,313-420 |  | pcDNA3.1+ |  |
| NgR1-C | 1-310 | 313-420 |  | pcDNA3.1+ |  |
| NgR1-AB | 1-310 | 313-420 |  | pcDNA3.1+ |  |
| *Domain deletion* |  |  |  |  |  |
| NgR1ΔNT | 1-25,58-473 | | | pcDNA3.1+ | Data not shown |
| NgR1Δ1-2 | 1-57,106-473 | | | pcDNA3.1+ | Data not shown |
| NgR1Δ3-4 | 1-105,155-473 | | | pcDNA3.1+ | Data not shown |
| NgR1Δ5-6 | 1-154,203-473 | | | pcDNA3.1+ | Data not shown |
| NgR1Δ7-8 | 1-202,251-473 | | | pcDNA3.1+ | Data not shown |
| NgR1ΔCT | 1-259,311-473 | | | pcDNA3.1+ | Data not shown |
| NgR1ΔGPI-CT | 1-310 | | | pcDNA3.1+ | Data not shown |

Sequences used from the following reference sequences:
\* NgR1: NM\_023004 (modified from OriGene SC126683)
\*\* NgR2: NM\_178570 (modified from OriGene SC310413)
\*\*\* NgR3: NM\_178568 (modified from OriGene RC216415)

### Preparation of purified proteins

#### Purified NgR1(1-447)-Fc Protein

Purified full-length NgR1 tagged with a human IgG (R&D Systems; 1208-NG) was resuspended in sterile PBS and stored at −80°C until use.

#### Purified NgR1(1-310)-Fc, JAM-A-Fc, and CAR-Fc Proteins

Plasmids encoding the extracellular domains of JAM-A and CAR fused C-terminally to a rabbit IgG Fc have been described [52]. Using sticky ended cloning PCR, JAM-A sequences were replaced with NgR1 a.a. 1-310 to engineer a NgR1(1-310)-Fc cDNA. Soluble NgR1(1-310)-Fc, JAM-A-Fc, and CAR-Fc proteins were produced in mammalian cells and purified by the Vanderbilt Antibody and Protein Resource using protein-A column affinity.

### Purified NgR1-His Protein

Using round-the-horn primers and PCR mutagenesis, sequences encoding a 7His-tag and an early stop codon were appended to the nucleotides correlating to a.a. 310 of NgR1. FreeStyle™ 293-F cells (Thermo Fisher Scientific) were transiently transfected at a cell density of 1 × 10^6^ cells/mL using a 1:3 ratio of plasmid DNA and polyethylenimine (PEI 25K™, Polysciences). Cells were incubated at 37°C for 7 days in a humidified incubator containing 8% CO_2_. Cells were pelleted at 7,000 rpm at 4°C for 15 min. Supernatants were collected and supplemented with cOmplete™ protease inhibitor EDTA free (Roche) per manufacturer’s directions and loaded onto a 5 mL HisTrap column (Cytvia) preequilibrated with buffer (10 mM HEPES pH 7.4, 150 mM NaCl, and 20 mM imidazole). A 20 – 500 mM imidazole gradient was added to the column to first wash and then elute NgR1-His. NgR1-His was further purified by size exclusion chromatography using a HiLoad 16/60 Superdex 200 column (Cytiva) and collected into buffer (10 mM HEPES, pH 7.4, and 150 mM NaCl). The protein was concentrated to 8 mg/mL and frozen at −80°C until use.

### Functionalization of AFM tips

AFM tips were functionalized with T1L, T3D, or T3SA+ virions using NHS-PEG_27_-aldehyde linkers as described [104]. Amino-functionalization of the tips was conducted using aminopropyl-triethoxysilane (APTES; Sigma-Aldrich) in gas phase [105]. Tips are immersed in chloroform (Sigma-Aldrich) for 5 min and then cleaned with UV radiation and ozone (UV-O; Jetlight) for 15 min. Tips were placed inside a desiccator containing two plastic trays and previously flooded with argon gas. APTES (30 µL) and triethylamine (10 µL) were pipetted separately into the trays and incubated for 2 h in the sealed chamber. Trays were removed and the desiccator was flooded with argon gas for 10 min. AFM tips were then left to “cure” the APTES coating for at least 2 days. Following amino-functionalization, AFM tips were coupled with flexible PEG linkers. First, the tips were immersed in a solution containing NHS-PEG_27_-aldehyde (3.3 mg) in chloroform (0.5 mL) and triethylamine (30 µL) for 2 h. Then, the tips were rinsed three times with chloroform for 5 min. After allowing the tips to dry, they were placed on Parafilm in a polystyrene Petri dish, which was stored in an ice box. Next, 100 µL of virus (10^9^ particles/mL) was pipetted onto the tips and 2 µL of freshly prepared sodium cyanoborohydride solution (∼6% [wt/vol] in 0.1 M NaOHaq) was added to the virus droplet. The tips were incubated in this solution at 4°C for 1 h, after which 5 µL of ethanolamine (1 M [pH 8.0]) was mixed into the droplet. Tips were incubated at 4°C for 10 min, rinsed 3X in ice-cold virus buffer (150 mM NaCl, 15 mM MgCl_2_, 10 mM Tris [pH 7.4]), and stored in virus buffer at 4°C until use.

### Preparation of NgR1-coated model surfaces

Soluble full-length NgR1 protein (R&D Systems, AF1208-NG) was immobilized on model surfaces using Protein A chemistry as described [106]. Gold-coated surfaces were rinsed with ethanol, dried with a low nitrogen flow, cleaned with UV radiation and ozone (UV-O; Jetlight) for 15 min, rinsed with ethanol, and dried with nitrogen gas. Surfaces were incubated overnight in an ethanol solution containing 1% COOH (1 mM 16-mercaptohexadecanoic acid) and 99% OH (11-mercapto-1-undecanol) alkanethiols. The next day, surfaces were rinsed with ethanol, dried with nitrogen gas, and incubated for 30 min in a solution of NHS and EDC (10 mg/mL NHS and 25 mg/mL 1-ethyl-3-[3-dimethylaminopropyl]-carbodiimide [EDC]). Surfaces were washed 3X with milliQ water and incubated at RT for 1 h in 100 µL of Protein A (100 µg/mL) in PBS. Surfaces were rinsed 3X with washing buffer (PBS containing 0.05% Tween20) and non-specific binding sites were blocked by incubating surfaces for 1 h in 300 µL of blocking buffer (PBS containing 0.05% Tween20 and 1% BSA). NgR1-Fc (50 µL of 100 µg/mL) was incubated on surfaces for 1 h. NgR1-coupled surfaces were rinsed with washing buffer and stored in PBS until use.

### Force-distance-based AFM on model surfaces

Force-distance curve-based AFM experiments on model surfaces were conducted in PBS at RT using virus-functionalized MSCT-D probes (spring constants were calculated using thermal tune [107], with values ranging from 0.030 to 0.047 N/m). Force-Robot300 (Bruker, Germany) and Nanoscope Multimode 8 (Bruker, Nanoscope software v9.1) atomic force microscopes, operated in the force volume (contact) mode were used to conduct these experiments. Gold-coated surfaces grafted with NgR1 were mounted on a piezoelectric scanner using a magnetic carrier. All experiments were conducted in PBS and areas of 5 ˣ 5 µm were scanned, with 32 ˣ 32 pixel resolution (corresponding to 1,024 force-distance curves) and a ramp size set to 500 nm. The approach velocity was kept constant at 1 µm/s and the maximum force was set to 500 pN.

Dynamic force spectroscopy (DFS) experiments to measure multiple loading rates were conducted by varying retraction speeds, set to 0.1, 0.2, 1.0, 5.0, 10, and 20 µm/s with no surface delay. Kinetic on-rate (k_on_) measurements were conducted by measuring the binding probability for hold times of 0, 50, 100, 150, 250, 500 and 1000 ms, allowing the tip to stay in contact with the surface for different periods of time.

In a separate set of experiments, as an independent negative control, surface blocking was conducted by incubated surfaces with anti-NgR1 antibody (R&D Systems; 100 µg/mL). Measurements were taken at a retraction velocity of 1 µm/s, before and after the addition of the antibody. The same sample area was probed several times, using the same tip, with no surface delay.

### Force-distance-based AFM and fluorescence microscopy on living cells

Force-distance, curve-based AFM experiments on living cells were conducted by acquiring correlative images with an inverted epifluorescence microscope (Zeiss Observer Z.1) coupled to an atomic force microscopy (AFMi) (Bioscope Resolve, Bruker), operated in the PeakForce QNM mode (Nanoscope software v9.2). The AFMi was equipped with a 150-µm piezoelectric scanner. A 40x oil objective (NA = 0.95) was used. Cells were maintained in growth medium and at 37°C. A gas mixture supplemented with 5% CO_2_ and 95% relative humidity was infused at 0.1 L/min into the chamber. PFQNM-LC cantilevers (Bruker) were oscillated at 0.25 kHz with an amplitude of 750 nm. The sensitivity of each cantilever was calculated through thermal noise method. Images were taken at 256 ˣ 256 pixels, probing an area of 25 - 30 µm, with imaging forces of 500 pN and a scan frequency of 0.125 Hz.

In a separate set of experiments, as an independent negative control, anti-NgR1 antibody (R&D Systems; 50 µg/mL) was added to the cells’ medium in order to probe specific interaction between the virus and the sample. All the experimental parameters were kept the same and data were collected before and after the antibody addition.

### AFM Data analysis

Data acquired in experiments on model surfaces was analyzed using JPK Data Processing (version 6.1.149) and Nanoscope analysis software (v2.0, Bruker). Peaks corresponding to adhesion events between virus particles and NgR1 were selected and retraction force-distance curves were fitted with the worm-like chain model for polymer extension [108]. Regarding DFS experiments, loading rates were determined using the slope of the force-time curves and rupture forces were extracted. Origin software (OriginLab) was used to graph DFS plots and to fit histograms of rupture force distributions, applying various force spectroscopy models, as described [47, 109]. For kinetic on-rate analysis, the binding probability (BP; fraction of curves that displayed a binding event) was determined for the different hold times (*t*; the time the tip is in contact with the surface). Data were fitted and K_D_ calculated as described [110].

For experiments on living cells, AFM images were analyzed with Nanoscope analysis software (v2.0, Bruker) and Gwyddion. ImageJ (v1.52e) was used to calculate the binding probability through pixel enumeration. Optical images were analyzed with Zen Blue software (Zeiss). Force-distance curves showing adhesion events were analyzed with NanoScope analysis software (v2.0, Bruker) and Origin software (OriginLab).

### Receptor cDNA transfection and infection of CHO cells

The day before transfection, 10^5^ CHO cells/well were seeded into 24-well tissue culture plates. Cells were transfected with cDNA using FuGene 6 (Promega, E2691) following manufacturer’s guidelines and a ratio of 0.5 µg of plasmid:1.5 µL FuGene 6 (Promega, E2691) in Opti-MEM (Gibco). “Mock”-transfected indicates that cells received only Opti-MEM. Cells were incubated at 37°C for 48 h post-transfection before infection or flow cytometric analyses.

For infections, transfected CHO cells were inoculated with T3SA-diluted in PBS^-/-^ at an MOI of 10 PFU/cell and incubated at 37°C for 1 h. Virus was removed and cells were incubated in completed Ham’s F12 medium. At 24 hpi, cells were washed with PBS^-/-^ and fixed with cold methanol at −20 °C for at least 30 min. Fixed cells were washed and incubated with reovirus polyclonal serum diluted in 0.5% Triton X-100 in PBS, followed by incubation with Alexa Fluor 488-labeled secondary IgG. Cells were counterstained with DAPI and imaged at 10X using a Lionheart FX automated imaging system (BioTek). Infected cells (FFU) were enumerated per field for at least four fields of view per well in triplicate wells.

### Flow cytometry assessment of reovirus binding or receptor expression

Monolayers of mock- or cDNA-transfected CHO cells were washed with PBS, detached using CellStripper Dissociation Reagent (Corning), and quenched with an equal volume of PBS^-/-^ supplemented to contain 2% FBS (FACS buffer). All further incubations, washes, and pelleting were conducted in racked titer tubes (Bio-Rad; 2239391) on ice or at 4°C. Washes and virus incubations were conducted in PBS^-/-^ and antibody incubations were conducted in FACS buffer. Cells were pelleted at 500 ˣ *g* at 4°C for 3 min, washed, and resuspended with T3SA- (10^5^ particles/cell) on ice for 1 h. Cells were pelleted, washed 3X with PBS^-/-^ to remove unbound virus, and incubated with reovirus-, NgR1-, mixed NgR1/NgR2-, or myc-specific antibodies at 4°C for 1 h. Cells were washed 3X, stained with Alexa-labelled secondary antibody at 4°C for 1 h, and washed again 3X. Cells were fixed in PBS^-/-^ supplemented to contain 1% paraformaldehyde and analyzed by flow cytometry. Results were quantified using FlowJo software.

### PI-PLC removal of NgR1 from the cell surface

At 48 h post-transfection, CHO cells were treated with 100 μL of either Opti-MEM (Gibco) or PI-PLC from Bacillus cereus (Invitrogen; LSP6466) resuspended to 0.5 U/mL in Opti-MEM (Gibco). Cells were incubated at 37°C for 1 h, washed in cold PBS^-/-^ and lysed in RIPA buffer prior to interrogation of protein expression by immunoblot analysis as described below.

### Expression and purification of the σ3_3_μ1_3_ heterohexamer

High Five™ insect cells (1 L) were infected with 20 mL of third passage recombinant baculovirus expressing μ1 and σ3 of reovirus strain T1L. Cells were incubated at 27°C for 72 h, pelleted by centrifugation at 500 ˣ *g* for 10 min, and resuspended in heterohexamer lysis buffer (150 mM KCl, 20 mM Tris [pH 8.5], 2 mM MgCl_2_, 2 mM β-mercaptoethanol [BME], 3 mM PMSF, and completed with benzonase [1,000 units] and cOmplete™ protease inhibitor cocktail [Roche] prior to use). Cells were lysed using an Avestin Emulsiflex and debris was pelleted at 31,200 rpm at 4°C for 40 min. The supernatant was loaded into a MonoQ ion-exchange column and washed with buffer A (20 mM Tris pH 8.5, 20 mM MgCl2, 2 mM BME) containing 150 mM NaCl. Heterohexamer was eluted with a linear buffer A gradient from 150 to 450 mM NaCl. Fractions containing μ1 and σ3 were pooled and concentrated to 1 mL with a 30 kDa molecular weight cut-off (MWCO) centricon. Protein was diluted to 4 mL with buffer A lacking NaCl and mixed dropwise with ammonium sulfate buffer (4 M ammonium sulfate, 20 mM Tris pH 8.5) to a final concentration of 0.7 M ammonium sulfate. Protein was loaded into a phenyl sepharose column and eluted with a linear buffer A gradient from 0.7 M to 0 M ammonium sulfate. Fractions containing heterohexamer were pooled and concentrated with a 30 kDa MWCO centricon. Protein was passed through a Superdex 200 size-exclusion column equilibrated with buffer B (20 mM Bicine pH 9, 100 mM NaCl, 2 mM MgCl2, 2 mM BME). Heterohexamer fractions were pooled and concentrated to 1 mg/mL.

### Production of 35S-labeled σ3

*In vitro* transcription and translation reactions were conducted using the TNT coupled rabbit reticulocyte lysate system (Promega, L4610) according to the manufacturer’s instructions. Reactions were supplemented with RNasin Plus RNase Inhibitor (N2611) and [^35^S]-methionine (PerkinElmer, NEG709A500UC). Plasmids encoding T1L or T3D S4 were added and reactions were incubated at 30°C for 1-1.5 h. Translation reactions were terminated with 3-fold excess of stop buffer (20 mM HEPES-KOH [pH 7.4], 100 mM potassium acetate, 5 mM magnesium acetate, 5 mM EDTA, 2 mM unlabeled methionine) supplemented prior to use with a final concentration of 1 mM dithiothreitol (DTT) and 2 mM puromycin.

### Immunoprecipitations, Native PAGE, SDS-PAGE, immunoblotting, and phosphor imaging

#### Immunoprecipitation

To immunoprecipitate proteins translated in vitro in RRLs, 5 µg of mouse 10C1 or 8H6 antibody or NgR1-Fc or CAR-Fc were incubated with σ3-expressing reactions at 4°C for 1 h with rotation. Samples were added to Protein G Dynabeads and incubated at 4°C for 1 h with rotation. The flow-through containing unbound protein was resolved by native PAGE. Dynabeads were washed four times with Tris-buffered saline (20 mM Tris-HCl pH 7.5, 150 mM NaCl) containing 0.1% Tween-20, eluted with SDS sample buffer, and resolved by SDS-PAGE.

#### Native PAGE

Samples for native PAGE were diluted in 4X Native PAGE Sample Buffer (ThermoFisher, BN2003) and loaded into wells of 4-16% Native PAGE Bis-Tris acrylamide gels (ThermoFisher). Samples were electrophoresed using the blue native PAGE Novex Bis-Tris gel system (ThermoFisher). Light blue anode buffer was supplemented with 0.1% L-cysteine and 1 mM ATP.

Electrophoresed proteins were transferred to PVDF. Membranes were soaked in 8% acetic acid for 15 min, rinsed with ddH2O, dried, incubated with 100% methanol for 1 min, rinsed with ddH2O, blocked with 5% bovine serum albumin (BSA) diluted in PBS^-/-^ and incubated with antibodies for immunoblotting. The NativeMark Protein Standard (ThermoFisher, LC0725) provided molecular weight estimation.

#### SDS-PAGE

Samples for SDS-PAGE were diluted in 5X SDS-PAGE sample buffer and incubated at 95°C for 10 min. Samples were loaded into wells of 10% acrylamide gels (BioRad, 4561036) and electrophoresed at 100V for 90 min. Following electrophoresis, proteins were either stained with colloidal blue (ThermoFisher, LC6025) or transferred to nitrocellulose for immunoblotting. Immunoblot analysis was conducted as described [111] and gels were scanned and analyzed using an Odyssey CLx imaging system (Li-Cor).

#### Phosphor Imaging

Following electrophoresis, gels were incubated in 40% methanol/10% acetic acid at RT for 1 h, washed three times with ddH2O, and dried on filter paper at 80°C for 2 h using a BioRad model 583 gel dyer. Dried gels were applied to a phosphor imaging screen for 12 - 36 h, followed by imaging using a Perkin Elmer Cyclone Phosphor System Scanner (B431200). Protein bands labeled with 35S-methionine were analyzed using ImageJ software (244).

### Cryo-EM data collection and processing

Purified T3SA-reovirus virions were incubated in PBS alone or with soluble NgR1-His (in a ratio of 1 σ3: 4 NgR1-His) at 4°C for 4 h. Quantifoil R 3.5/1 copper grids were glow discharged and 4 µL sample was plunge-frozen in liquid ethane using a Vitrobot Mark IV (Thermo Fisher). Frozen hydrated specimens were imaged at 300 kV using low-electron-dose conditions on a JEOL 3200 FSC cryo-electron microscope equipped with a direction detector K2 summit camera (Gatan) in the NIH-funded National Center for Macromolecular Imaging at Baylor College of Medicine. Movie stacks were collected semi-automatically using SerialEM [112] at 20,000x magnification, corresponding to a pixel size of 1.71 Å. Each image stack was fractioned in 50 subframes with a total dose of 0.55 electrons/A². Image stacks were aligned and beam induced motion was corrected using the *alignframes* package from IMOD [113]. Contrast transfer function (CTF) correction was conducted using *e2ctf_auto.py* from EMAN2.3 [114, 115]. 7,182 and 11,046 particles were initially picked using *e2.boxer.py* for the reovirus and reovirus:NgR1 complex sample, respectively, and reference-free two-dimensional (2D) class averages were computed using *e2refine2d.py*. Good class averages with clear viral features were selected to calculate an initial three-dimensional (3D) model using *e2initialmodel.py* with applied icosahedral symmetry. The initial model was low-pass filtered to ∼30 Å and used as model for 3D-refinement using *e2refine_easy.py*. After several rounds of refinement, 5,954 and 1,648 high-quality particles were selected using *e2.evalrefine.*py for the reovirus and reovirus:NgR1 sample, respectively. 3D reconstructions were refined to a final resolution of 7.2 Å and 8.9 Å, respectively, according to the Gold Standard Resolution FSC plot (threshold 0.143) of EMAN.

### Model fitting and analysis

The 3D reconstructed maps allowed fitting of protein coordinates using the *Fit in map* function of UCSF Chimera [116]. The following atomic coordinates from the Protein Data Bank (PDB) were used: PDB-ID *1ozn* for NgR1 [54], *1jmu* for the μ1σ3 heterohexamer [64], and *3iyl* for the core proteins λ2, σ2, and λ1 [117]. To minimize clashes between the fitted protein chains, a single round of *phenix.real_space_refine*, including simulated annealing, was conducted [118]. The difference density map was calculated using the *vop* command in UCSF Chimera. RMSD values were calculated using the *align* and *rms_cur* algorithms in PyMol [119]. Figures were prepared using UCSF Chimera [116] and PyMol [119].

### Statistical analysis

Statistical tests were conducted using Prism 7 (GraphPad Software) and Origin (OriginLab). Means of individual experiments are shown for experiments conducted fewer than five times. *P* values of less than 0.05 were considered to be statistically significant. Descriptions of the specific tests used are provided in the figure legends.

## ACKNOWLEDGMENTS

We thank Laurie Silva for critical review of an early version of this manuscript. This work was supported by the U.S. Public Health Service awards R01 AI038296 (D.M.S. and T.S.D.) and R01 AI118887 (D.M.S., T.S., and T.S.D.). Additional support was provided by UPMC Children’s Hospital of Pittsburgh (P.A.), the Heinz Endowments (T.S.D.). Support (M.K., R.S., and D.A) was also contributed by the Universiteì catholique de Louvain, the Fonds National de la Recherche Scientifique (F.R.S.-FNRS), the European Research Council by the European Union’s Horizon 2020 research and innovation program (grant number 758224), and the FNRS-Welbio (grant number CR-2019S-01). M.K. (Postdoctoral Researcher) and D.A. (Research Associate) are members of the FNRS. The funders had no role in study design, data collection and analysis, decision to publish, or preparation of the manuscript.

## SUPPLEMENTARY FIGURE LEGENDS

**Appendix Figure S1.**
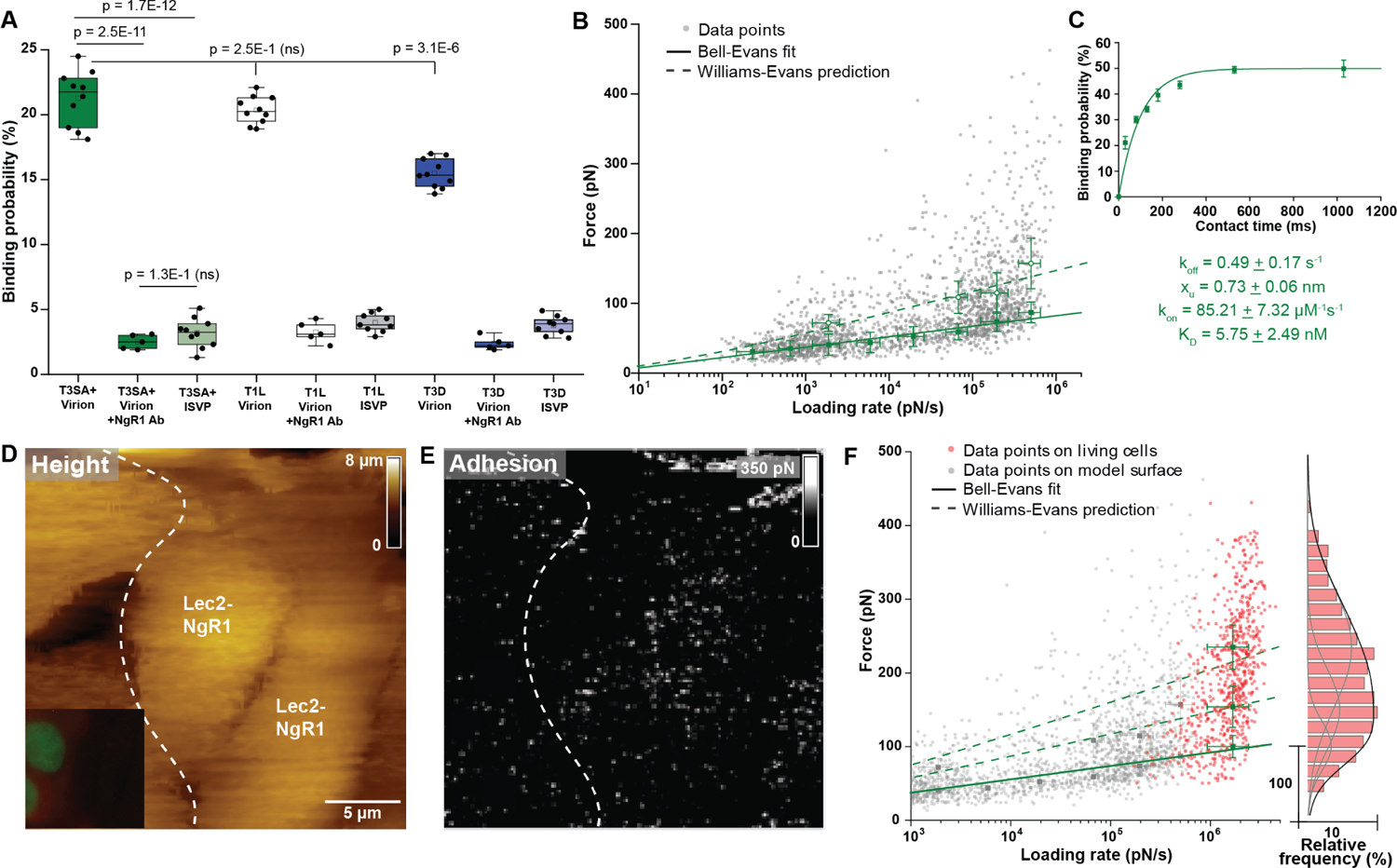
Reovirus T3SA+ virion binding to NgR1 on model surfaces and living cells resembles T1L more closely than T3D. A Box plot summary showing the probability of T3SA+ binding to NgR1 (green) relative to that of T1L (white) and T3D (green), before and after injection of NgR1-specific antibody. One data point represents the binding probability from one map acquired at a retraction speed of 1 µm/s. The square in the box indicates the mean, the colored box the 25th and 75th percentiles, and the whiskers the highest and lowest values. The line in the box indicates the median. N = 10 maps examined for 3 independent experiments. *P* values were determined by two-sample *t*-test. B, C Dynamic force spectroscopy (B) and contact time (C) analysis of T3SA+ binding to an NgR1 model surface, yielding the k_off_, *x*_u_, k_on_, and K_D_ values of this interaction. D, E, F Binding of T3SA+ virions to NgR1 probed on receptor-overexpressing living cells (Lec2-NgR1). Topography (D) and adhesion (E) maps of the probed area shown in the insert. DFS plot of T3SA+ virion interactions with NgR1 on model surfaces (grey dots, from panel B) and living cells (red dots). Histogram of the force distribution observed on cells fitted with a multi-peak Gaussian distribution (N > 800 data points) is shown on the side. Error bars indicate SD of the mean values. Data are representative of at least N = 10 cells from 3 independent experiments.

**Appendix Figure S2.**
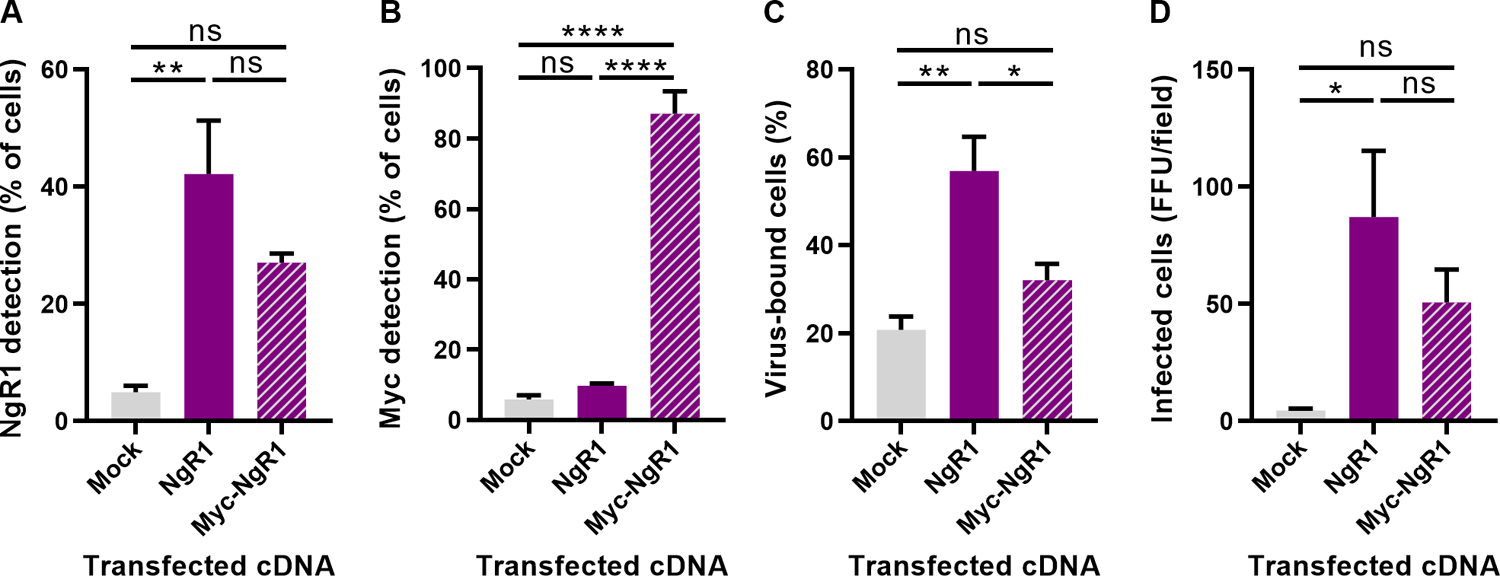
N-terminally tagged NgR1 is expressed well but permits diminished reovirus binding and infectivity. A, B, C, D CHO cells were mock-transfected or transfected with the cDNAs shown and incubated for 48 h. (A) NgR1 expression, (B) Myc expression, or (C) reovirus binding were determined by flow cytometry. (D) Infectivity was assessed by FFU assay. Error bars indicate SD of the mean. Values that differ significantly from mock by one-way ANOVA and Dunnett’s test are indicated (*, *P* < 0.05; **, *P* < 0.01; ****, *P* < 0.001). ns = not statistically significant.

**Appendix Figure S3.**
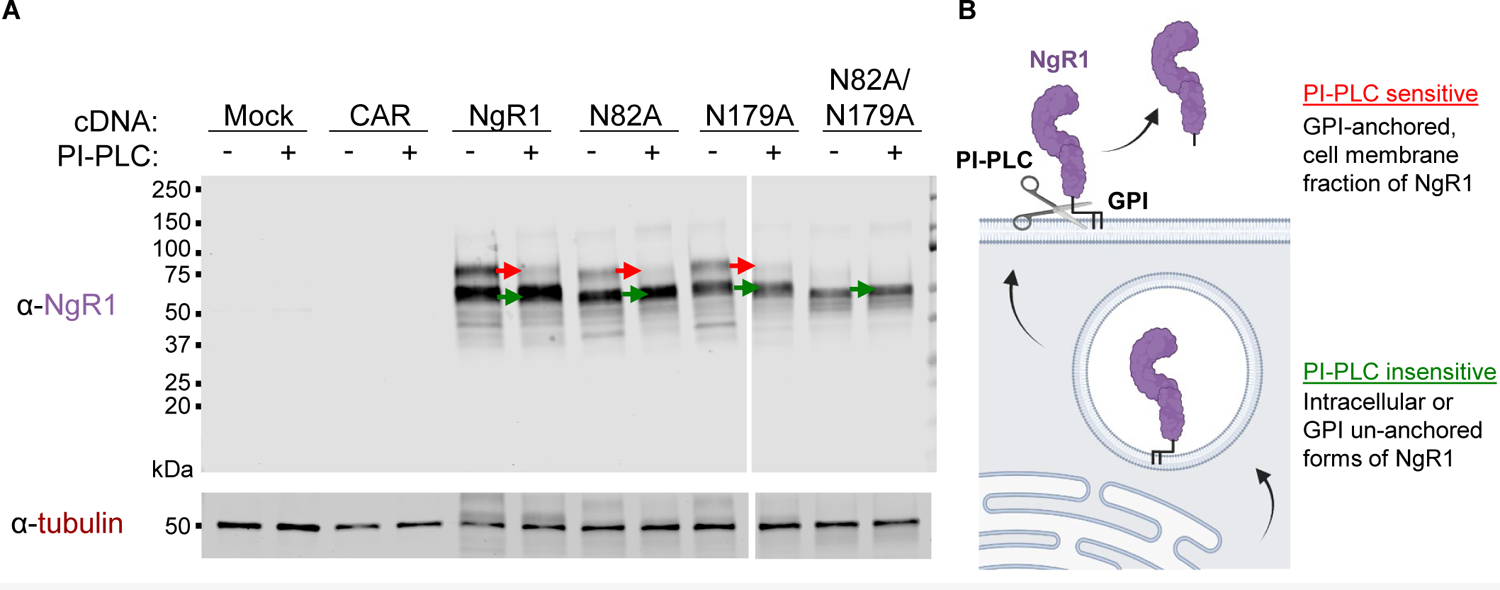
Slower-migrating NgR1 bands in polyacrylamide gels are sensitive to phospholipase treatment. (A) CHO cells were mock-transfected or transfected with the cDNAs shown, incubated for 48 h. Cells were treated with vehicle control or phosphatidylinositol phospholipase C (PI-PLC), washed, and lysed in a detergent buffer. Lysed cell extracts were resolved by SDS-PAGE and probed using either NgR1-specific anti-sera or a monoclonal antibody recognizing tubulin. (B) PI-PLC is membrane impermeable and cleaves GPI-anchored proteins from the cell surface. Schematic made using BioRender.

**Appendix Figure S4.**
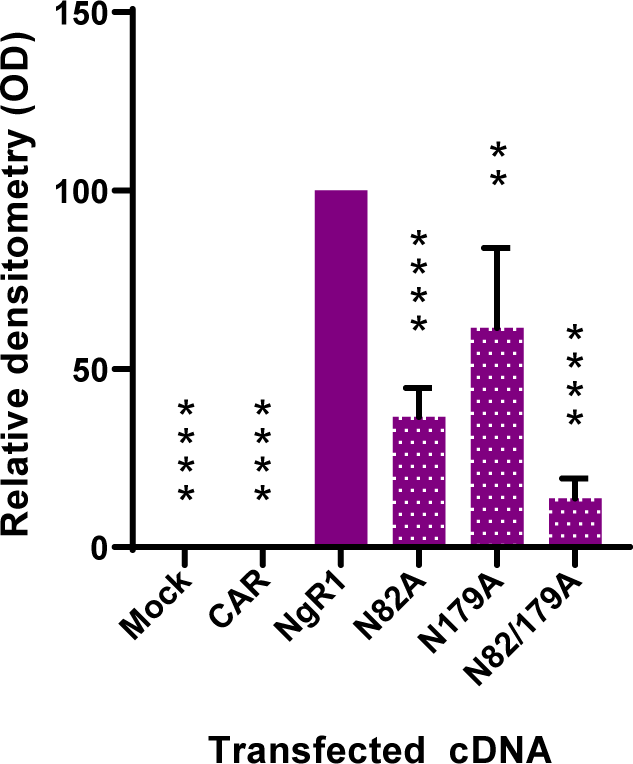
Quantification of NgR1 detected in cell lysates by immunoblot analyses. CHO cells were mock-transfected or transfected with the indicated cDNAs and incubated for 48 h. Cells were lysed and membrane-cleared fractions were electrophoresed by SDS-PAGE and transferred to nitrocellulose. Membranes were probed for either NgR1 or tubulin and imaged (Fig 3J). Quantification relative to background and tubulin of 3 independent gels is shown.

**Appendix Figure S5.**
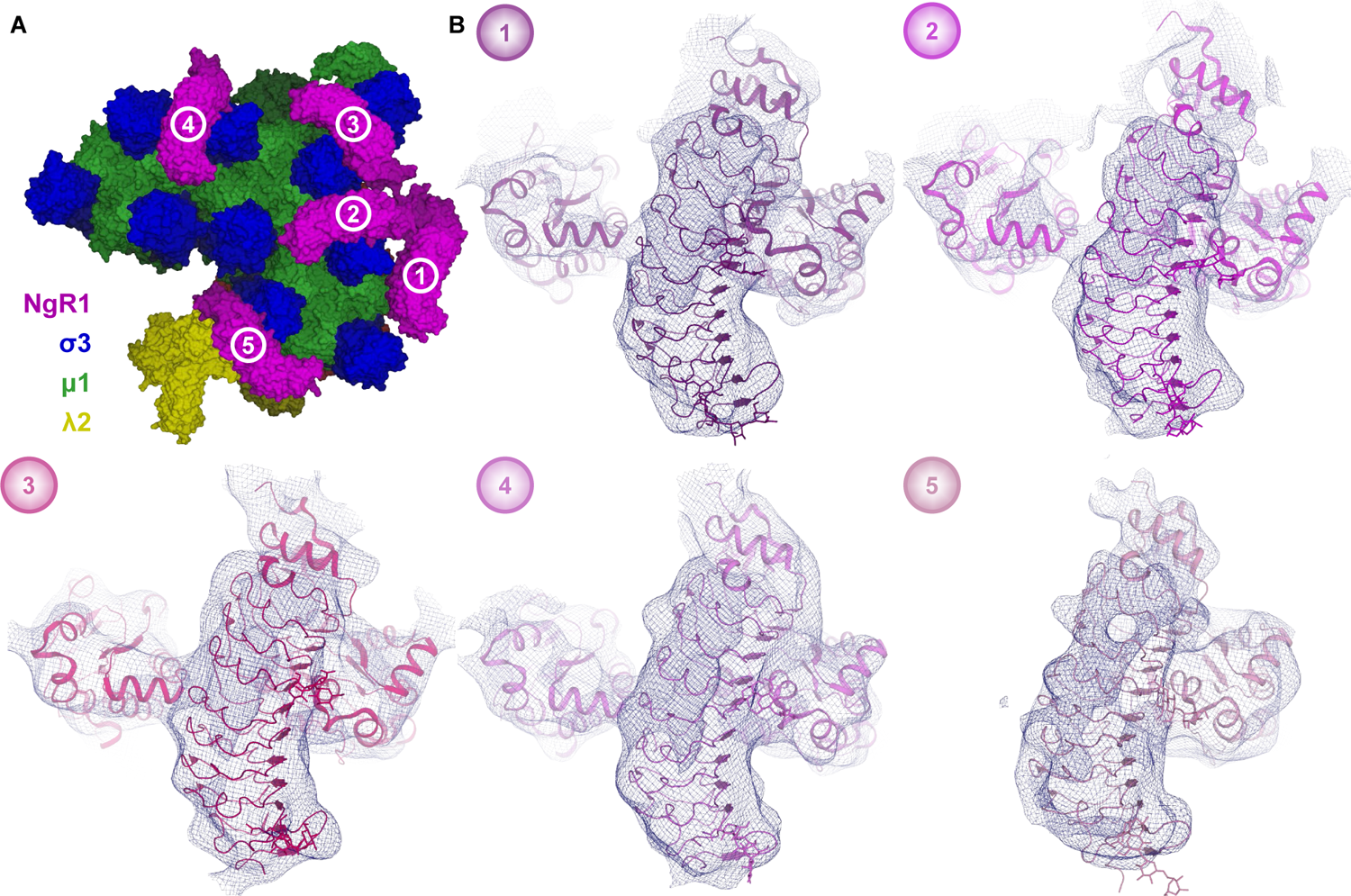
NgR1 molecules bind at different sites on the reovirus virion surface with similar conformations. A Surface representation of the modelled asymmetric unit of a reovirus virion in complex with NgR1, with NgR1 molecules labelled 1-5 (extracted from Figure 5 molecular docking). B Binding mode comparison of all NgR1 molecules within the asymmetric unit. The corresponding 3D reconstruction at counter level 1.0 is shown as blue meshes. For NgR1-(1), the σ3 protomer to the right was included from a σ3_3_ µ1_3_ heterohexamer from the neighboring asymmetric unit.

